# Parkinson’s disease uncovers an underlying sensitivity of subthalamic nucleus neurons to beta-frequency cortical input

**DOI:** 10.1101/513234

**Authors:** Magdalena K. Baaske, Eszter Kormann, Abbey B. Holt, Alessandro Gulberti, Colin G. McNamara, Monika Pötter-Nerger, Manfred Westphal, Andreas K. Engel, Wolfgang Hamel, Peter Brown, Christian K.E. Moll, Andrew Sharott

**Affiliations:** MRC Brain Network Dynamics Unit, Department of Pharmacology, University of Oxford, Oxford OX1 3TH, United Kingdom; Nuffield Department of Clinical Neurosciences, John Radcliffe Hospital, University of Oxford, Oxford, UK; Department of Neurophysiology and Pathophysiology, University Medical Center Hamburg-Eppendorf, 20246 Hamburg, Germany; Department of Neurology, University Medical Center Hamburg-Eppendorf, 20246 Hamburg, Germany; Department of Neurosurgery, University Medical Center Hamburg-Eppendorf, 20246 Hamburg, Germany

**Keywords:** Cortex, beta oscillation, Parkinson’s disease, STN neuron, synchronisation

## Abstract

Abnormally sustained beta-frequency synchronisation between the motor cortex and subthalamic nucleus (STN) is associated with motor symptoms in Parkinson’s disease (PD). It is currently unclear whether STN neurons have a preference for beta-frequency input (12-35Hz), rather than cortical input at other frequencies, and how such a preference would arise following dopamine depletion. To address this question, we combined analysis of cortical and STN recordings from awake PD patients undergoing deep brain stimulation surgery with recordings of identified STN neurons in anaesthetised rats. In PD patients, we demonstrate that a subset of STN neurons are strongly and selectively sensitive to fluctuations of cortical beta oscillations over time, linearly increasing their phase-locking strength with respect to full range of instantaneous amplitude. In rats, we probed the frequency response of STN neurons more precisely, by recording spikes evoked by short bursts of cortical stimulation with variable frequency (4-40Hz) and constant amplitude. In both healthy and dopamine-depleted animals, only beta-frequency stimulation selectively led to a progressive reduction in the variability of spike timing through the stimulation train. We hypothesize, that abnormal activation of the indirect pathway, via dopamine depletion and/or cortical stimulation, could trigger an underlying sensitivity of the STN microcircuit to beta-frequency input.

## Introduction

Abnormally synchronised oscillations at beta frequencies (12-35Hz) are robust features of the cortico-basal ganglia network in patients with Parkinson’s disease (PD) (Brittain JS et al. 2014). These activities are reduced by successful treatment of akinesia and rigidity, either with dopaminergic medication or deep brain stimulation (DBS) (Marsden JF et al. 2001; Levy R et al. 2002; Williams D et al. 2002; Kuhn AA et al. 2008). Moreover, this enhanced neuronal synchronisation can be replicated in rodent and primate models of PD (Bergman H et al. 1994; Sharott A et al. 2005). Whether or not beta oscillations are causal in generating akinetic motor symptoms, a question still under debate, they provide a useful biomarker for akinetic/rigid symptoms of the disease (Zaidel A et al. 2009; Stein E and I Bar-Gad 2012; Brittain JS and P Brown 2014). This is evident in the multiple studies that have found positive correlations between metrics of beta oscillations and symptom severity (Kuhn AA et al. 2005; Kuhn AA et al. 2008; Kühn AA et al. 2009; Little S et al. 2012; Sharott A et al. 2014; Neumann WJ et al. 2016; Sharott A et al. 2018) and the utilisation of their temporal dynamics to drive effective closed-loop DBS (Little S et al. 2013).

The subthalamic nucleus (STN) has proved a particularly important network node in studying properties of pathological oscillations in PD. In humans, the majority of studies have recorded oscillations either within the STN, or between cortex and STN. While investigations in experimental animals have clearly demonstrated these activities extend across the entire network (Mallet N, A Pogosyan, LF Marton, et al. 2008; Pasquereau B and RS Turner 2011; Brazhnik E et al. 2016; Deffains M et al. 2016; Sharott A et al. 2017), the STN likely has a pivotal role (Deffains M et al. 2016). This stems, in part, from its position as the only group of excitatory neurons within the basal ganglia network. STN neurons lock to beta-frequency population oscillations in cortex, measured by ECoG and EEG, and local field potentials (LFPs) in the STN and other basal ganglia structures (Mallet N, A Pogosyan, LF Marton, et al. 2008; Moran A et al. 2008; Tachibana Y et al. 2011; Shimamoto SA et al. 2013; Sharott A et al. 2018). Several studies have demonstrated that cortical oscillations lead those in the STN (Williams D et al. 2002; Fogelson N et al. 2006; Lalo E et al. 2008; Sharott A et al. 2018), suggesting that STN neurons are entrained by cortical beta oscillations. It remains to be elucidated whether this entrainment is due to intrinsic resonance, network dynamics or a combination of the two.

On the one hand, like all basal ganglia neurons outside the striatum, STN neurons fire autonomously via a specific set of intrinsic mechanisms (Nakanishi H et al. 1987; Bevan MD and CJ Wilson 1999). On the other hand, the STN is integrated in the subcortico-cortical microcircuit with specific network dynamics. The most prominent inputs to STN neurons are excitatory synapses from pyramidal tract neurons in the frontal cortex, often referred to as the “hyperdirect” pathway (Nambu A et al. 2002), and inhibitory synapses from the globus pallidus external segment (GPe) (Smith Y et al. 1998; Bevan MD et al. 2002). Input from the GPe can also be heavily influenced by cortical output through the cortically-driven striatal indirect pathway neurons (Kita H and T Kita 2011; Zold CL et al. 2012). When isolated from their inputs, the autonomous firing of STN neurons consists of highly regular single spiking (Bevan MD and CJ Wilson 1999). Irregular firing, bursting and oscillation are thus the result of the synaptic inputs perturbing this resting firing pattern (Wilson CJ 2015). Such external perturbations in firing pattern are dependent on the relative timing of excitatory and inhibitory inputs relative to the intrinsic currents driving autonomous firing (Wilson CJ 2013, 2015). Despite the convergence of both excitatory and inhibitory afferents on single neurons and rich oscillatory input from cortex (Murthy VN and EE Fetz 1992, 1996), STN neurons do not display sustained oscillations or synchrony in the healthy brain (Bergman H et al. 1994; Wilson CJ 2013; Deffains M et al. 2016). A key question, therefore, is, whether the mechanism to synchronise preferentially with beta-frequency cortical input is always present, but is prevented by some kind of active decorrelation that is lost in PD (Wilson CJ 2013).

Here, we demonstrate that STN neurons display selective phase-locking and temporal dynamics relative to the instantaneous amplitude of network oscillations, specific to the beta-frequency band. In PD patients, we find that the spike-timing of STN neurons is more influenced by input in the beta-frequency range, even when lower frequency oscillations have higher spectral power. Using recordings from anaesthetised rats, we manipulated the frequency of cortical input to identified STN neurons, while keeping the amplitude constant. These experiments suggested that beta-frequency input is optimal for reducing the variance in STN spike timing in both control and parkinsonian animals. Together, our findings suggest that STN neurons are selectively vulnerable to beta-frequency input under physiological, as well as pathological conditions.

## Methods

### Human Data

#### Patient information

To address the question how firing of STN neurons is related to ongoing population oscillations in humans we analysed neuronal activity from patients undergoing deep brain stimulation surgery targeting the STN. The study was conducted in the agreement with the Code of Ethics of the World Medical Association (Declaration of Helsinki, 1967) and was approved by the local ethical committee. All patients gave their written informed consent to participate in the study. All patients included in the study suffered from advanced Parkinson’s disease and did not show cognitive impairment as reflected by the Mattis Dementia Rating Scale (clinical details are given in supplementary Table 1). Dopamine agonists were replaced by levodopa >7 days and levodopa was paused on the day of surgery. The operation was conducted under local anaesthesia and systemic analgosedation with remifentanile (0.01-0.1 μg/kg/min) and paused during the procedure of microelectrode recordings. The detailed surgical procedure is described elsewhere (Hamel W et al. 2003; Moll CK et al. 2014; Sharott A et al. 2014). The stereotactic coordinates for the STN were 11-13 mm lateral to the midline, 1-3 mm inferior and 1-3 mm posterior to the mid-commissural point on both sides. The rostral and caudal borders of the STN were determined by electrophysiological criteria (i.e. increase in background noise, irregular tonic discharge pattern, oscillatory and burst patterns) (Sterio D et al. 2002; Sharott A et al. 2014).

#### Data acquisition and recording setup

STN LFP and unit recordings were collected during the standard neuro-navigation procedure applied for clinical electrode placement. Microelectrode signals were performed in the BenGun configuration (MicroGuide and NeuroOmega, Alpha Omega, Nazareth, Israel) with 1-3 parallel electrodes. Each electrode was 2 mm from the central tip, which was aimed at the final target position. Microelectrode signals were amplified (x20.000), band-pass filtered (300-6000 Hz) and digitized (sampling rate: 20kHz or 44kHz). Local field potentials obtained from the macro-tip 3mm above the micro-tip (contact size ≈ 1mm, impedance <1kΩ) were band-pass filtered between 1 and 300 Hz, amplified (x5000-10000) and sampled at 1375 or 3005 Hz. All signals were referenced to the uninsolated distal part of the guide tube. Simultaneous cortical recordings were made using either the electrocorticogram (ECoG) from the dura (approximately above dorsolateral prefrontal cortex) or the EEG positioned at Fz according to the international 10-20 system, both re-referenced to Pz. LFPs were not recorded in two patients, which were therefore only included to analysis of cortical signals and units.

#### Data selection and processing

Recordings had a variable length (109 +/− 77 s), and depending on recording quality, total recordings time and trajectories, the number of extracted single and multi-units varied between patients (18.88 +/− 9.67 units per patient). Spike detection and spike sorting was performed offline using Offline Sorter (Plexon Inc., Dallas, TX, USA). The threshold was set for each individual recording based on the distribution relative to the background noise, typically >4 SD. Single units were than extracted by a manual sorting procedure in 2D and 3D feature space based on several waveform parameters as principal components (signal energy, peak time and the presence of a trough in the auto-correlogram) (Sharott A et al. 2014; Sharott A et al. 2018). Units corresponding to the putative spiking activity of up to 3 neurons and thus not fulfilling the criteria for well isolated single unit activity were classified as MUA, but were clearly distinct from background activity (Sharott A et al. 2018). The mean firing rate of included units was 37.62 Hz +/− 18.84 Hz (n=302). Note that units classified as MUA (n=173) had a slightly higher firing rate (39.72 +/− 20.24 Hz) than SUA (n=129, 34.84 +/− 16.43 Hz) (MWUT, *P*=0.04). Only stationary units with a stable firing rate >5 Hz over 3 second bins for at least 30s were included in analysis. Note that while both SUA and MUA were used in analysis to increase n numbers, similar results could be observed when only SUA was used. All signals were down-sampled to a common sampling rate of 1000 Hz prior to all analyses.

#### Spectral analysis

Prior to spectral analysis all signals were low-pass filtered at 200 Hz and notch-filtered at multiples of 50 Hz using digital zero-phase forward and backward filtering. Power spectra were calculated using a multi-taper approach with 3.5 slepian tapers and a frequency resolution of 0.5 Hz. Power spectra were normalized for 1/f by using a fitted pink noise spectrum for each recording (Feingold J et al. 2015).

Oscillations in spike-trains were detected by calculating the compensated power spectral density. The power spectral density was calculated using a hanning window with a frequency resolution of 0.49 Hz (NFFT 2048) and then normalized by a spectrum from 100 shuffled spike-trains (Rivlin-Etzion M et al. 2006). Confidence limits were calculated by using the compensated power in the range from 300 to 500 Hz. Significant peaks were detected at a significance level p<0.05 which was corrected for the number of frequency bins in the range of interest between 0-100 Hz (Bonferroni corrected alpha significance level 0.05/206 bins: p<2.43e-04). For comparison with the phase-locking analysis we calculated the mean compensated power of the preferred frequency of phase-locking +/− 5 Hz and normalized it by the power between 300-500 Hz.

#### Phase-locking Analysis

To determine the relationship between unit activity and ongoing oscillations detected in field potential recordings (EEG, ECoG, LFP) we performed a phase-locking analysis. The field signal was filtered in exponentially increasing frequency bands from 0-100 Hz using a 2^nd^ order Butterworth filter. The instantaneous phase of the oscillation was extracted for each spike using the Hilbert transform, and the distribution of spike-field phases compared to a uniform distribution was assessed using the Rayleigh test (Berens P 2009). Significant locking was defined by a Rayleigh test with p<0.05. We then calculated the mean vector length as a measurement of phase-locking strength (Lachaux JP et al. 1999) using circular statistics (Berens P 2009). In this formalism the vector length is a measurement of the variance of instantaneous spike-field phases and can reach a value between 0 and 1. To control for the effect of differing number of spikes, (Vinck M et al. 2010; Sharott A et al. 2012) a z-score of vector length was calculated using 100 ISI shuffled spike trains. A z-score over 2 was defined to represent significant phase-locking.

#### Magnitude dependent phase-locking

To determine whether STN unit phase-locking was dependent on oscillation magnitude, the phase-locking analysis was repeated for spikes occurring at select magnitudes of the field oscillation. The absolute values of the Hilbert transform were used to extract the instantaneous magnitude of a given oscillation. The distribution of spike-time magnitudes was divided into deciles (i.e. the 1^st^ decile is corresponding to the magnitude from 0 to the border of decile 1, (i.e. 1^st^ decile is from 0-10% of the spike-time magnitude distribution, each part contains 1/10 of the magnitude distribution values, schematic illustration in Fig.2A). The magnitude distribution was created based on magnitude values only occurring at spike times which lead to equal numbers of spikes in each decile, thus the calculation of phase-locking strength in each decile is based on equal number of spikes. In this formalism, calculated phase-locking strength at low magnitude values of the ongoing oscillation correspond to the 1^st^ decile, with increasing magnitude values up to the highest values at the 10^th^ decile.

We first analysed every available pair in exponentially increasing frequency bands between 0 and 100 Hz. To gain a deeper understanding what happens selectively within the beta-frequency range we repeated the analysis for significantly phase-locked units (Rayleigh test p<0.05) in a single beta-frequency band. Because peak beta frequencies could differ between patients and possibly also units, we selected the centre frequency between 12 and 40 Hz with the lowest Rayleigh statistic p-value for each recording. To determine the lowest decile, where a unit starts significant phase-locking, the first decile, when the z-score exceeded a value of 2 and stayed high until the 10^th^ decile, was detected.

#### Correlation analysis

We calculated Pearson’s correlation coefficient between the vector length and the normalised magnitude at each decile for each filter frequency. Values with a Cook’s distance > 1 were excluded prior to the correlation analysis (Cook RD 1977). For characterizing the shape of the correlation selectively in the beta-frequency range we repeated the correlation analysis with the mean magnitude in each decile, normalised by the mean magnitude of the highest decile. In case of the phase-locking analysis to LFPs, the vector length was averaged for each unit to compensate for multiple LFP channel combinations with a given unit. We performed this analysis for each recording and averaged across the group.

#### Beta burst analysis

To analyse phase-locking during beta bursts, field signals (LFP, ECoG, EEG) were band-pass filtered using 11 overlapping frequency bins from 10-40 Hz (5Hz wide, 2.5 Hz overlap, 2^nd^ order Butterworth filter with digital zero phase forward and backward filtering). For each STN-unit – field pair, the filtered frequency with the lowest p-value of the Rayleigh statistic was chosen. We then applied the previously described clinically relevant threshold of the 75^th^ percentile of the magnitude of the filtered signal and detected continuous episodes of elevated beta power above that threshold, referred to as beta bursts (Tinkhauser G, A Pogosyan, S Little, et al. 2017; Tinkhauser G, A Pogosyan, H Tan, et al. 2017; Tinkhauser G et al. 2018). Only beta bursts with a minimum duration of 3 cycle lengths were included in the analysis. We than repeated the phase-locking analysis of all the spikes occurring within 1 cycle wide bins before, during, and after the beta burst. A minimum of 20 spikes within all summed cycle bins corresponding to one position relative to the burst was required for calculation of vector length. In addition, we calculated a Pearson’s correlation between the deciles of magnitude and the mean magnitude in each decile and performed statistics on a group of STN-unit-LFP/ECoG/LFP pairs, which followed the magnitude and were positive correlated and a group, which was uncorrelated.

#### Rate analysis

The mean firing rate in each magnitude decile and in each bin relative to the beta burst was calculated to determine if increases in phase-locking strength were related to rate changes.

### Rat Data

#### Juxtacellular recordings of STN neurons and cortical stimulation in control and 6-OHDA lesioned rats

Experiments were carried out on 5 healthy control male rats and 9 6-OHDA hemi-lesioned male rats (adult Sprague-Dawley rats, Charles River) (190-200g) under general anaesthesia and were conducted in accordance with the Animals Act (Scientific Procedures, 1986, United Kingdom) and the Society for Neuroscience Policies on the Use of Animals in Neuroscience Research.

#### Induction of unilateral 6-hydroxydopamine (6-OHDA) hemi-lesions

Unilateral hemi-lesions of dopaminergic neuron in the substantia nigra pars compacta were induced as described previously (Magill PJ et al. 2001). Briefly, animals were anesthetised with isoflurane (4% v/v isoflurane O2), and 1 μl (6mg/ml) of 6-OHDA-toxin solution was injected over 10 min with a glass pipette (diameter 18 μm) through a small burr hole either in the medial forebrain bundle (4.1 mm posterior of bregma, 1.4 mm lateral to the midline, 7.9 mm ventral to the dura) (2 rats) or in the substantia nigra pars compacta (4.5 mm posterior of bregma, 1.2 mm lateral to the midline and 7.9 mm ventral to the dura) (7 rats). 25 minutes prior to this desipramine (25 mg/kg, Sigma, Poole, UK) was injected i.p. to protect noradrenergic neurons from cell death. Electrophysiological recordings were conducted at least 2 weeks or later post lesioning. Successful lesions were confirmed on the same sections by a significant reduction in Thyrosine-hydroxylase (TH) immune-reactivity in the striatum of the lesioned hemisphere in comparison to the intact side.

#### Electrophysiological recordings and juxtacellular labelling

Electrophysiological recordings in the STN were performed under general anaesthesia with induction by isoflurane (4% v/v isoflurane in O2) and were maintained with ketamine (30 mg/kg i.p., Willows Francis) and xylazine (3 mg/kg, i.p. Bayer). Body temperature, heart rate and peripheral reflexes were monitored throughout. Single unit activity and LFPs were recorded using a glass electrode (tip diameter 1.5-2.5 μm, impedance measured in situ 10-30 MΩ) filled with 0.5 M NaCl solution and Neurobiotin (1.5% w/v, Vector Laboratories) targeting the STN (Magill PJ et al. 2004). Signals were analogue amplified (x10, Axoprobe-1A amplifier (Molecular Devices), further amplified (x100, NL-106 AC-DC Amp, Digitimer) and sampled at 16.6 kHz using a Power 1401 Analag-Digital converter and Spike 2 acquisition and analysis software (Version 7.2, Cambridge Electronic Design). Unit activity was band-pass filtered between 300-5000 Hz (DPA-2Fs filter/amplifier, Digitimer) and LFPs were low-pass filtered at 2000 Hz (NL125 filters, Digitimer). For verification of exact recording positions a subset of putative STN-neurons were juxtacellularly labelled (Pinault D 1996; Magill PJ et al. 2000) with Neurobiotin. During this procedure positive current pulses (2-10 nA, 200 ms, 50% duty cycle) were applied until the neuron’s spike train showed entrainment by the current. The location was then post hoc confirmed by localising the Neurobiotin filled cell bodies within the borders of STN using standard immunohistochemistry methods. The position of unlabelled neurons was reconstructed based on the position of labelled neurons recorded in the same animal (Nakamura KC et al. 2014).

#### Electrical stimulation of the motor cortex

For cortical stimulation parallel, bipolar stimulating electrodes (constructed from nylon-coated stainless steel wires, California Fine Wire, tip diameter 100 μm, tip separation 150 μm, impedance 10 kΩ) were implanted in the primary motor cortex (M1, stereotactic coordinates: 4.2 mm anterior and 3.5 mm lateral to Bregma, (Watson C and G Paxinos 1986)) ipsilateral to the nigrostriatal lesion (Magill PJ et al. 2004). The depth of the cortical electrode (2.2 mm below the dura) corresponded approximately to the layer 5/6 of M1. Stimulation was performed in 12 blocks of 5 stimuli at 4, 10, 15, 20, 25, 30 and 40 Hz with a 0.3 ms square-wave pulse (constant-current isolator, A360D, World Precision Instruments). Stimulation amplitudes ranged from 75 to 1000 μA, adjusted depending on strength of modulation.

#### Histology and imaging

Standard indirect immunofluorescence protocols were used to verify the recording position of Neurobiotin labelled somata in the STN and the success of dopaminergic terminal loss in the striatum (Magill PJ et al. 2001; Mallet N et al. 2012). Rats were given a lethal overdose of ketamine (150 mg/kg) and were transcardially perfused with 300 ml of 0.1% w/v glutaraldehyde and 4% w/v paraformaldehyde in 0.1 M phosphate buffer. Extracted brains were stored overnight in fixative solution at 4°C. 50 μm. Coronal sections were used for staining and washed 3 × 10 min in Triton PBS (PBS containing 0.3% v/v Triton X-100, Sigma). For visualization of STN borders we performed an immune-staining for the marker forkhead box protein P2 (FoxP2) (Hontanilla B et al. 1997; Campbell P et al. 2009). Sections were first overnight incubated with Cy3-conjugated streptavidin (1:1000 dilution, Life technologies) and then incubated for 1 hour in Triton PBS containing 10% v/v donkey serum (NDS, Jackson ImmunoResearch Laboratories) followed by incubation over night at 4°C with the primary anti-body (Triton PBS containing 1% v/v NDS with goat anti-FoxP2 antibody 1:500, Santa Cruz). After washing in Triton PBS, sections were incubated with the secondary antibody Alexa 488 flurophor (raised in donkey, 1:1000, Life Technologies). Then the washed, mounted and cover-slipped (Vectashield, Vector Laboratories) sections were inspected with an epifluorescence microscope (Zeiss Acio Imager M.2) for final confirmation of recording positions.

For verification of successful dopaminergic lesions coronal sections anterior-posterior +/− 0.5 mm from bregma were stained with tyrosine hydroxylase as marker for dopaminergic terminals and neurons. For indirect immunofluorescence the same incubation protocol as described above was used. Tyrosine hydroxylase immunoreactions (primary antibody: Chicken anti-TH, 1:500 Abcam; secondary antibody: Alexa Fluor 488 1:1000, Life Technologies) were carried out in 3 or more striatal and at least 1 nigral section. Only brains with a clear difference in the brightness of the lesioned and intact striatum were classified as successful hemi-lesioned animals.

#### Imaging

High resolution images of the confirmed STN neurons were taken using a confocal microscope (Zeiss LSM 880) equipped with 20x 0.8NAPlan-Apochromat and 40x 1.4NA Plan-Apochromat oil immersion objectives and Stereoinvestigator v11.0 software (MBF Biosciences). To capture the streptavidin and the FoxP2 immuno-reactivity the following lasers and filter-sets were used: Argon 488nm laser for Alexafluor 488 (emission detection: 493-554 nm) and HeNe 543nm for Cy3 (emission detection: 554-681 nm). Imaging was performed with 10x and 40x objectives. At 40x, a series of multiple-plane z-stacked images (optical section thickness 0.8 *μ*m) were captured in order to show the exact location of the somatodendritic structure of the well-filled STN neurons. Labelled STN neurons were visualised with FoxP2 by showing images of 63x magnification using 1.46NA objective, optimum x,y sampling according to Nyquist, 135 × 135 × 17 micron field of view/stack height, maximum intensity projection of 18 z planes (total height 17 microns). Images were opened with Fiji software (Schindelin et al., 2012). Both channels of the low magnification (20x) image were processed using ‘Enhance contrast’ (a linear contrast adjustment that automatically sets white and black points) set to 0.1% of pixels being saturated. The high magnification Z-stack images were flattened using standard deviation projection. Contrast were then enhanced with 0.8% saturation levels for the streptavidin channel.

The extent of DA lesion was evaluated using Zeiss Axio Imager M.2, equipped with Hamamatsu Flash 4.0 LT camera and Colibri 7 LED illumination. For assessing the extent of dopamine lesion, the whole coronal section was captured (+/−0.5 mm bregma) using Stereoinvestigator v11.0 software (MBF Biosciences) ‘Virtual Tissue’ scanning mode. Images were taken on a low magnification using a Plan-Apo 5x objective (numerical aperture 0.16). To image the TH signal, the AlexaFluor-488 channel was used (LED excitation at 475 nm, filter-set: 38). Once the whole section scans were performed, images were exported to Image J software without any further image processing. Then, a grid of 1000 × 1000 *μ*m was randomly overlaid on the image of the whole section and the most dorsolateral square (top corner of both side’s striatum) was chosen for measuring the brightness of the lesioned and non-lesioned hemispheres. In the same way, matching control areas were chosen from the cortex above each side. Mean brightness levels were measured for all four squares. Each striatal brightness measures were then divided by the corresponding cortical brightness measure and were compared with each animal. Animals were considered lesioned if there was more than 50% difference between the normalised brightness levels of the lesioned and non-lesioned striatum.

#### Peri-stimulus histograms of STN single unit activity

Peri-stimulus histograms (PSTHs) were calculated for each neuron for each frequency with summation across the course of stimuli with a bin size of 1 ms in the inter-stimulus interval. A z-score was calculated by subtracting the mean of the bin count in the pre-stimulus period of 500 ms before the occurrence of the first stimulus and dividing it by its standard deviation. Inclusion criteria for significant responses were a peak with a z-score above 2 and at least 6 trials with a spike after at least 4 out of 5 stimuli. These criteria had to be fulfilled for at least 5 frequencies. To achieve a significant response during the experiment, neurons often had stimulated with multiple amplitudes and the threshold of response could slightly vary between experimental sessions. Recordings leading to a significant response in the PSTH had a mean stimulation amplitude of 288 +/− 141 μA in controls and 281 +/− 153 μA in 6-OHDA lesioned, both in a range from 100-500 μA). Effects at different frequencies were compared within each neuron and across stimuli within each frequency.

#### Coefficient of Variation (CV) of the first spike following stimulus pulse across trials

For determining the variability of the intervals *(i)* between first spike and stimulus across the consecutive stimuli, we calculated the Coefficient of Variation *(CV)* across trials. The CV can reach a value between 0 and 1 and is defined as *CV=std(i)/mean(i)*. For the calculation of the CV for the first stimulus (because here there is no history to define stimulus frequency) all intervals from all frequencies were merged to get a more reliable measure for comparison.

#### Evoked LFP response

LFPs from rat recordings were obtained from the same electrode tip as single unit activity. For obtaining a spike-free LFP, spikes had to be removed from the LFP signal by thresholding the high-pass filtered signal at 3 standard deviations from the median of the stimulus artefact free signal. A window 1 ms before to 3 ms after a spike was replaced with a linear interpolation and a spike- and stimulus-free segment of data. Each recording was inspected visually, and the threshold was redefined if needed. To control for slight differences in stimulation amplitude between non-lesioned and lesioned animals, we selected amplitude matched unique recordings (n=12) from each group (1 recording of 160μA in controls was matched with 150μA in lesioned animals, all other pairs were exactly matched). For statistical comparison between experimental conditions we averaged the mean amplitude in each recording in the period from 5-15ms after the stimulus for estimating the amplitude of the peak.

#### Statistical analysis

Due to non-normal distribution of the data we used the Mann-Whitney-U-Test (MWUT) for pairwise comparison, unless otherwise stated. When more than 2 groups were available we performed a Kruskal-Wallis ANOVA with post hoc Dunn’s test unless otherwise stated. To account for within neuron variability, comparisons across consecutive stimuli were performed either by the signed Wilcoxon ranked test (2 comparisons) or the Friedman test (>3 comparisons) with post hoc Tukey-Kramer’s tests. All analyses were performed in MATLAB^®^ R2017a (Mathworks). Boxplots for visualization purposes were created in Graphpad Prism (GraphPad Software). Mean values +/− standard deviation and sample sizes are given in the result section. Error bars in all figures show SEM, boxplots show median with the 5-95 percentile, outliers not shown for overview reasons.

## Results

The aim of the study was to investigate the selectivity of STN neurons to beta-frequency input and the precise nature of any frequency preference. To address these questions, we utilised two approaches. Firstly, we investigated phase-locking dynamics of STN units to local field potentials and cortical field potentials from 16 PD patients undergoing DBS-surgery. Secondly, we applied short bursts of electrical stimulation to the motor cortex at different frequencies while recording in the STN of anesthetised 5 healthy and 9 dopamine-depleted rats.

### I. Entrainment of STN neurons to cortical and local oscillatory input in humans

#### STN neurons are more selectively phase-locked to beta oscillations than higher power sub-beta oscillations

Power spectra of LFP/ECoG recordings in patients with Parkinson’s disease have peaks in sub-beta, beta and gamma frequencies (Kuhn AA et al. 2005; Kempf F et al. 2009; Zaidel A et al. 2010; Shimamoto SA et al. 2013; West T et al. 2016). As LFP recordings are assumed to reflect the summarized synaptic input to the STN (Buzsaki G et al. 2012), we first investigated whether the power of these local and cortical oscillations is reflected in the activity/level of synchronisation of single STN neurons. While beta oscillations are implicated in PD motor symptoms, patients display higher power oscillations in the sub-beta (4-12 Hz) range (Fz (n=280, F(2,837)=308 *P*=1.31e-67, post hoc Dunn’s *P*<0.0001 between sub-beta/beta and sub-beta/gamma range, for beta/gamma range *P*=0.13), ECoG (n=186, F(2,555)=435.06 *P*=3.38e-95, post hoc Dunn’s *P*<0.0001 for all comparisons) and LFP (n=248, F(2,741)=417.49 *P*=2.20e-91, post hoc Dunn’s *P*=0.002 for comparison sub-beta/beta and *P*<0.0001 for beta/gamma, sub-beta/gamma), Figure 1A). In contrast, STN unit phase-locking strength was significantly higher to beta-frequency oscillations than sub-beta frequencies (Fz (F(2,837)=88.58 *P*=5.82e-20, post hoc Dunn’s *P*<0.0001 between sub-beta/beta and beta/gamma range), ECoG (F(2,555)=77.16 *P*=2.89e-17, post hoc Dunn’s *P*<0.0001 for all groups) and LFP (F(2,741)=60.95 *P*=5.81e-14, post hoc Dunn’s *P*<0.0001 between sub-beta/beta and beta/gamma range), Fig, 1B-D). Furthermore, units showed significantly stronger locking to beta-frequency activity than gamma (60-80 Hz). Therefore, in line with previous studies STN spiking was selectively responsive to beta frequencies, despite, higher alpha-frequency power (Moran A et al. 2008; Shimamoto SA et al. 2013; Sharott A et al. 2018).

**Figure 1.**
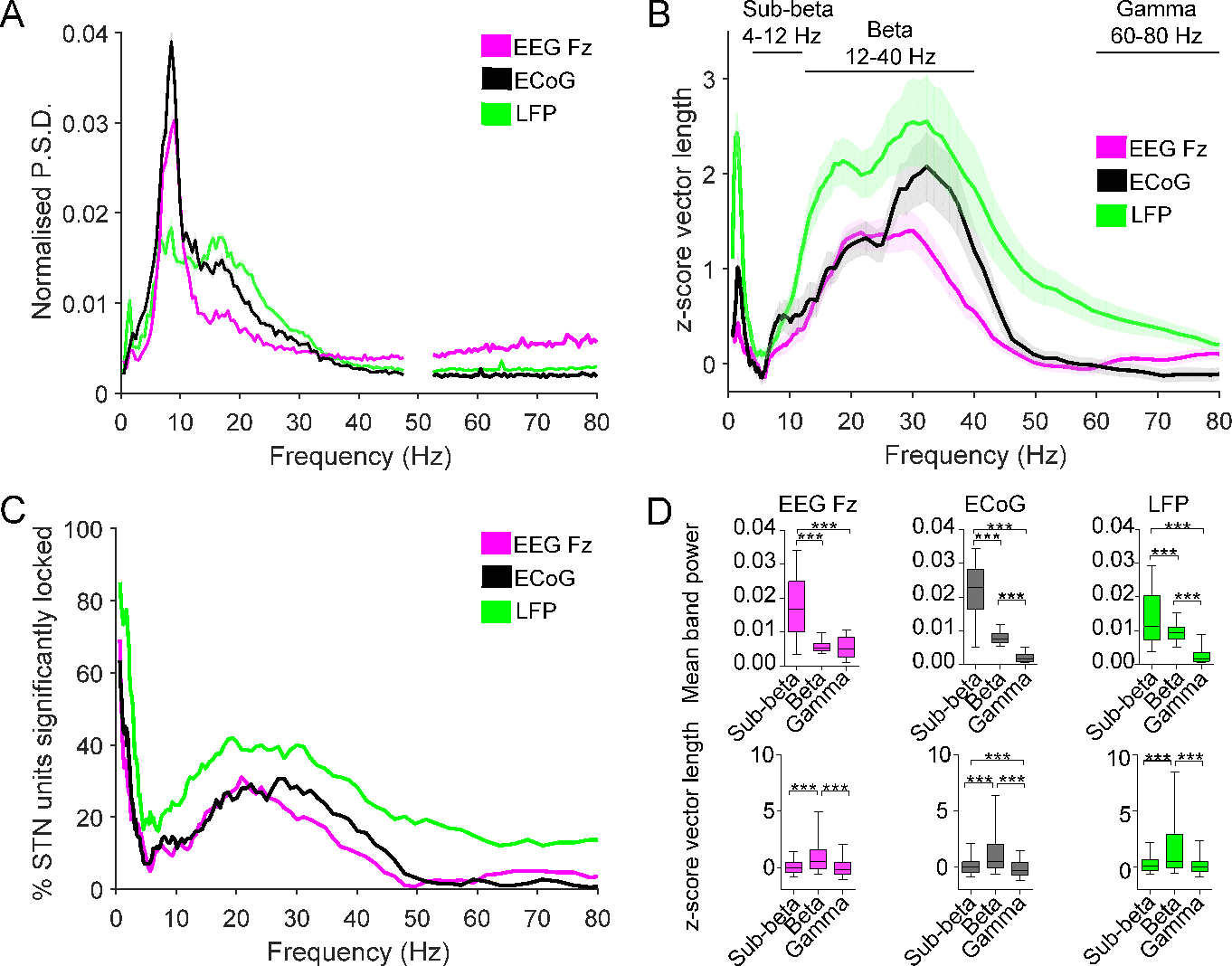
STN units were selectively phase-locked to beta oscillations despite the presence of higher power of oscillatory activity in the sub-beta-frequency range. **(A)** Mean 1/f normalized relative power spectral estimates normalized by the power from 0.5-80 Hz for EEG Fz (n=280), ECoG (n=186) and LFP (n=296). Shaded areas show +/− SEM. **(B)** Phase-locking strength for each of the STN units to EEG Fz (n=280), ECoG (n=186) and LFP (n=296). Note the selectively high phase-locking strength within the beta-frequency range. **(C)** The histogram of the proportion of significant phase-locked units, determined by a significant Rayleigh test, shows a broad peak in the beta-frequency range. **(D)** The mean relative power in sub-beta- (4-12 Hz), beta- (12-40 Hz) and gamma-(60-80 Hz) frequency range (top) and comparison of the phase-locking strength in the same frequency bands (bottom). Note the significant higher phase-locking strength of STN units to beta oscillations despite the higher spectral power for sub-beta oscillations. Significance was tested using a Kruskal-Wallis Anova with post hoc Dunn’s test. *** p<0.001. Box plots show the quartile bounderies with whiskers showing the 5-95 percentile.

#### Phase-locking strength of STN neurons to network beta oscillations was dependent on oscillation magnitude

Having established that STN neurons selectively lock to network oscillations in the beta-frequency range, we tested whether the strength of this synchronisation is predicted by the instantaneous magnitude of the oscillations. Spike-time oscillation magnitudes were divided into deciles (i.e. 1^st^ decile is from 0-10% of the spike-time magnitude distribution) (Fig. 2).

**Figure 2.**
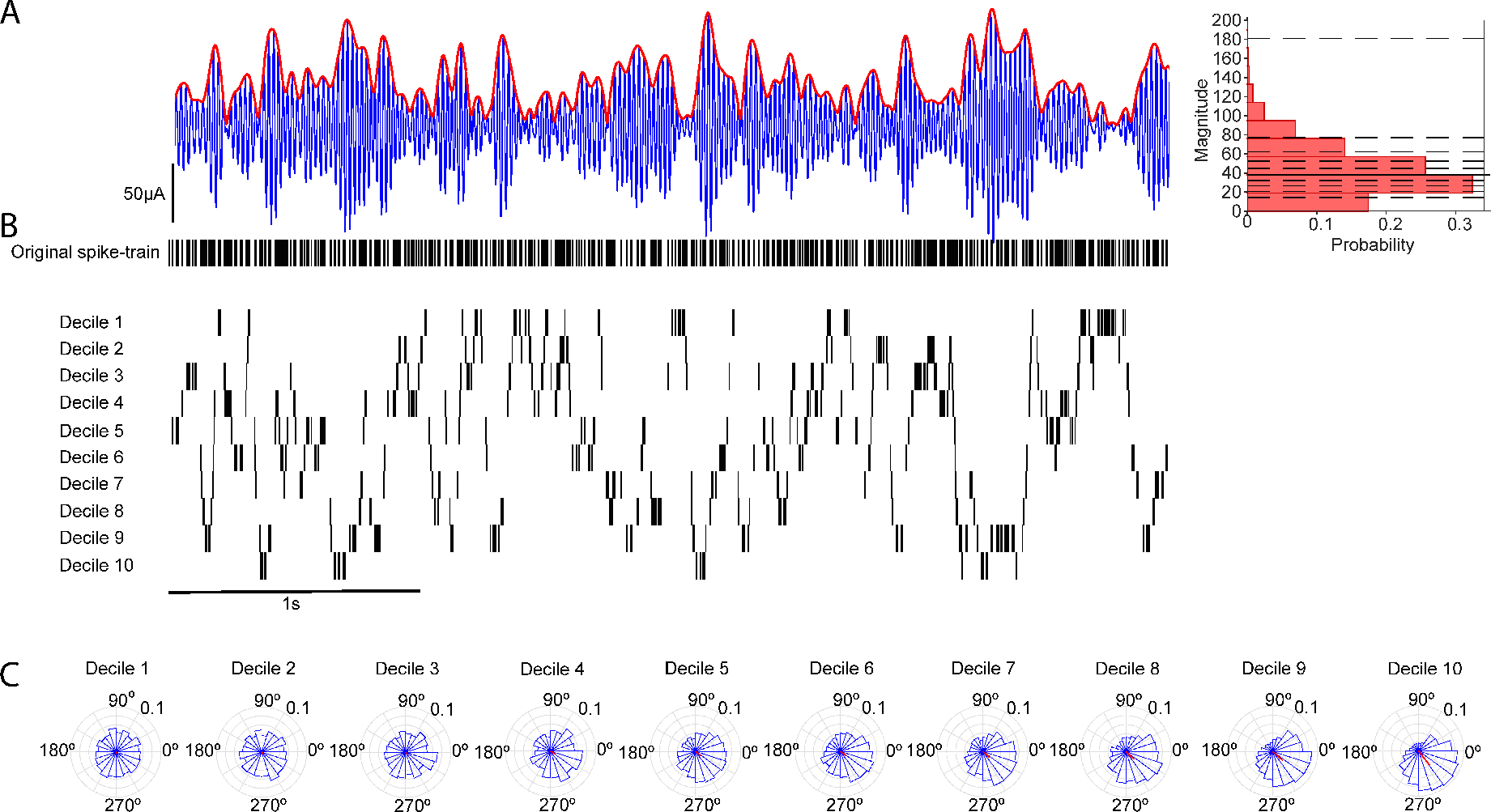
Magnitude dependent phase-locking analysis - schematic illustration. **(A)** Left, Example filtered LFP trace in the beta-frequency range (25-35 Hz) (blue) with instantaneous magnitude of the hilbert envelope (red). Right, Example distribution of spike-time magnitudes for the example recording shown in A (solid black line: median). Dashed black lines indicate decile divisions. **(B)** Original spike train (top) and spike trains generated by separating spikes occuring within each decile. Data corresponds with the LFP oscillation shown in (A). **(C)** Circular plots show the distribution of spike phases relative to the LFP beta oscillation for each decile. The red line shows the mean phase and vector sum. The vector length becomes longer and the spike phase distribution becomes more peaked with increasing deciles (magnitudes of the beta oscillation).

We were then able to calculate phase-locking values for spikes occurring within each decile, all with an equal number of spikes due to the magnitude distribution of magnitude values at spike times. For all investigated network oscillations (LFP, ECoG and EEG Fz), phase-locking strength of STN units increased with the magnitude of network beta oscillations, starting around the 6^th^/7^th^ decile (Fig. 3A,D,G). This correlation between oscillation magnitude and unit phase-locking strength was strongest in the beta-frequency range (Fig. 3B, E, H) (Comparison of the Pearson’s correlation coefficient between decile and z-score vector length: Fz (F(2,837)=85.90 *P*=2.23e-19, ECoG (F(2,555)=56.79 *P*=4.66e-13) and LFP (F(2,741)=63.13 *P*=1.96e-14), post hoc Dunn’s *P*<0.0001 for all comparisons).

**Figure 3.**
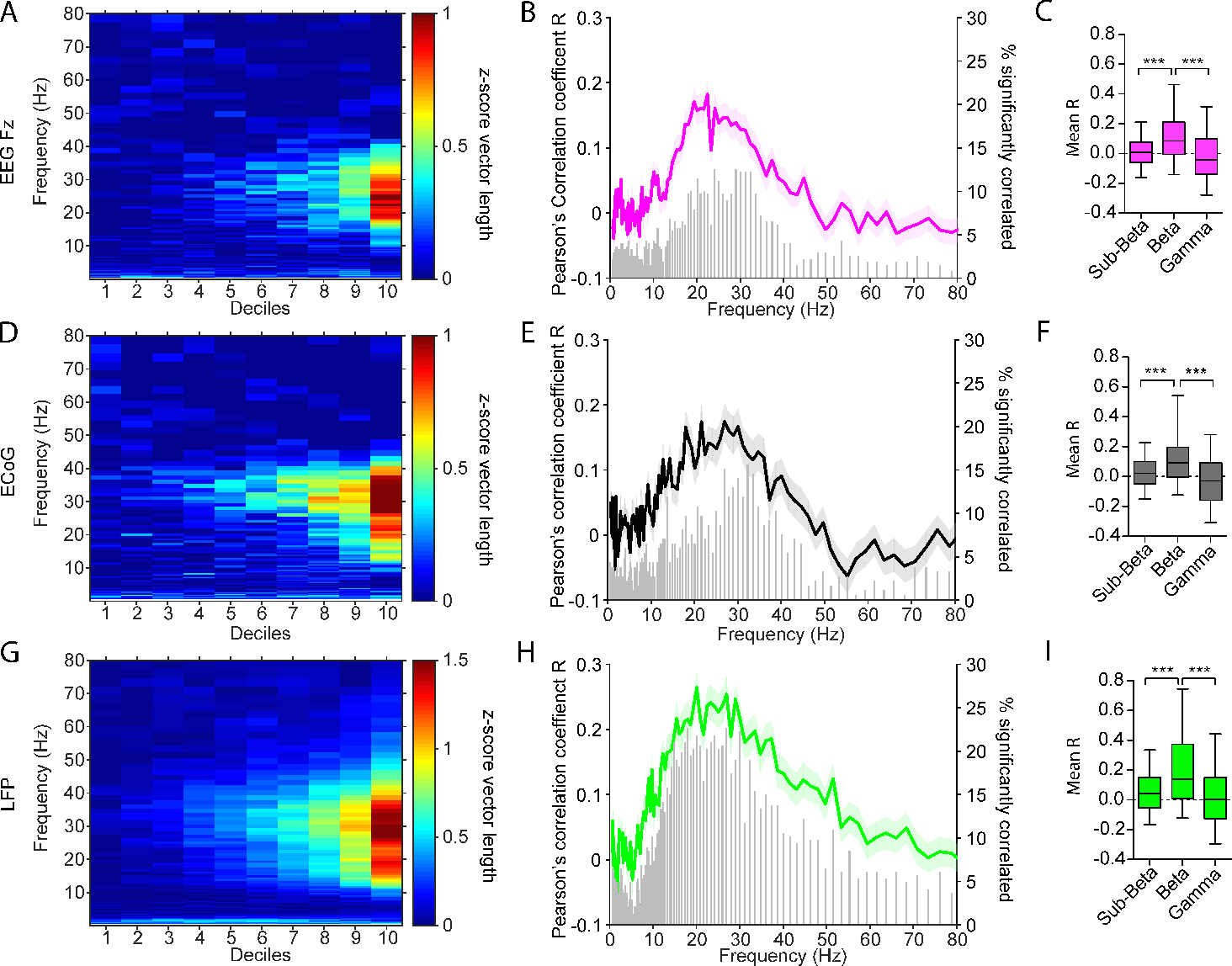
Magnitude dependent increase of STN unit phase-locking strength to network beta oscillations. **(A)** Mean Increase of phase-locking strength of STN units to Fz (n=280) in each decile for increasing frequency bins from 0-80 Hz. **(B)** For each recording and frequency the Pearson’s corellation coefficient (R value) between phase-locking strength and decile number (coloured trace, mean ± SEM, left y-axis) along with the number of significant correlated bins shown in the histogram (gray, right y-axis) were calculated. **(C)** Box plots show the mean correlation coefficient (between decile and z-score vector length) for the sub-beta-(4-12 Hz), beta-(12-40 Hz) and gamma-(60-80 Hz) frequency range (Kruskal-Wallis Anova with post hoc Dunn’s tests). **(D-F)** Show the same analysis of STN units with ECoG (n=186) and **(G-I)** with LFP (n=296) recordings. Note the magnitude dependent increase of phase-locking strength for all signals selectively in the beta-frequency range (A,D,G). The highest correlation and amount of bins significantly correlated was likewise highest in the beta-frequency range (B,E,H). The correlation strength was significantly higher in the beta-frequency range in comparison to sub-beta and gamma band power for all network population signals (C,F,I). *** p<0.001, whiskers of box plots show the 5-95^th^ percentile.

#### Entrainment of STN spiking to beta oscillations increases linearly with the instantaneous magnitude

The nature of the relationship between magnitude and synchrony could reveal the underlying transfer function describing how individual neurons are recruited into network oscillations. Because the magnitude of network oscillations is not normally distributed (Fig. 2A), but increased exponentially over the deciles, we repeated the correlation analysis with the mean normalised magnitude value in each decile (instead of decile number) to get a more accurate estimate of the shape of this relationship. Between 18-34% of significantly phase-locked STN units showed a significant positive correlation over the deciles with the magnitude of network oscillations (Fz n=28/154 (18 %), ECoG n=27/109 (25%) and LFP n=56/172 (33%)). The correlation of the mean vector length across deciles gave correlation coefficients close to 1 for all investigated population signals (Fig. 4 A-C, Fz R=0.96, *P*=3.35e-05; ECoG R=0.98, *P*=1.23e-06; LFP R=1, *P*=2.67e-09). To test whether the increase of phase-locking strength was dependent on firing rate, the same analysis was repeated for the firing rate at each decile. The overall firing rate in the majority of cases was not related to the phase-locking strength on a single recording basis (non-correlated firing rate-magnitude units: Fz: n=23/28 (82%), ECoG: n=20/27 (74%) and LFP: n=30/56 (54%)). Out of significantly correlated units, the majority were negatively correlated (Fz: n=5/28 (18%), EcoG: n=6/27 (22%) and LFP: n=16/56 (29%), with few significantly positively correlated (Fz: n=0/28 (0%), EcoG: n=1/27 (4%) and LFP: n=10/56 (18%)). The group average revealed a significant negative correlation for all network population signals (Fz n=28, Pearson’s R=-0.73, *P*=0.016; ECoG n=27, Pearson’s R=-0.78, *P*=0.007; LFP n=56, Pearson’s R=-0.66, *P*=0.03; Figure 4D-F). Neurons with the highest positive correlation coefficients between phase-locking strength and population amplitude also had the highest overall phase-locking values (Fz n=154, Pearson’s R=0.57, *P*=4.71e-15; ECoG n=109, Pearson’s R=0.55, *P*=8.67e-10; LFP n=172, Pearson’s R=0.51, *P*=6.99e-13; Fig. 4G-I). In addition, the mean phase-locking strength of positively magnitude-correlated units was significantly higher than that of uncorrelated units (MWUT, Fz *P*=7.16e-10; ECoG *P*=1.999e-08; LFP *P*=9.57e-21, Fig.4J-L).

**Figure 4.**
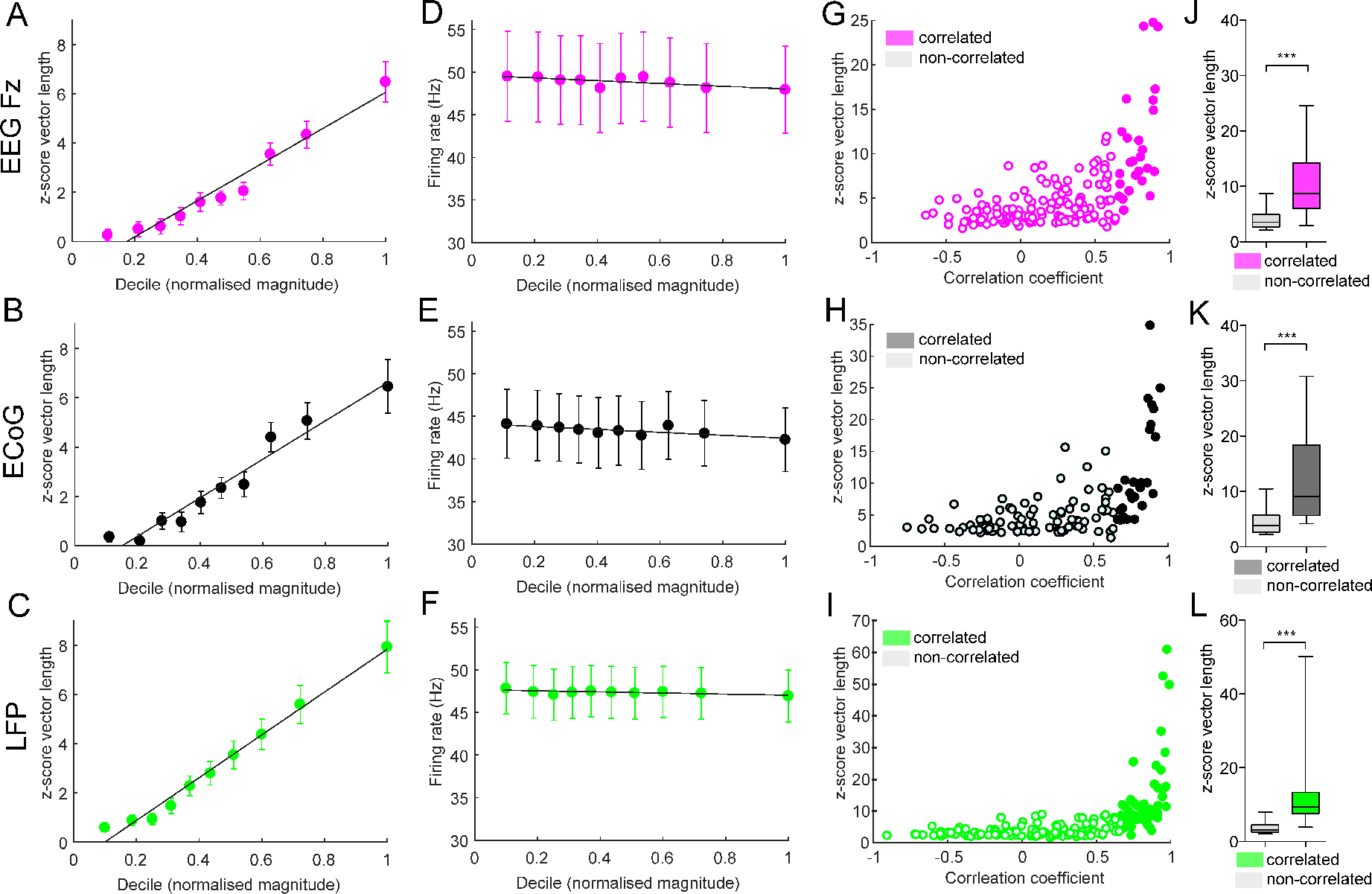
Linear entrainment of STN units to network beta oscillations. **(A,B,C)** Group average of phase-locking strength (z-score vector length) along the mean normalised magnitude in each decile (x-axis). Error bars show +/− SEM. Group means consist of positively correlated unit-network oscillation pairs only (i.e. units which phase-locking strength follows the magnitude of the network oscillation, Fz n=28/154 (A), ECoG n=27/109 (B) and LFP n=56/172 (C)). The line shows the linear fit of the group average with a significant correlation for all network oscillation signals (Fz R=0.96, *P*=3.35e-05; ECoG R=0.98, *P*=1.23e-06; LFP R=1, *P*=2.67e-09). **(D,E,F)** Group average of the firing rate +/− SEM in each decile along the mean normalized magnitude in each decile (x-axis) for Fz (D), EcoG (E) and LFP (F). Group averages were limited to consist of the same positively correlated unit-network oscillation pairs as in A-C. Note that the majority of units show no correlation between the magnitude deciles and the firing rate (see text) and the mean firing rate remained stable across the deciles. **(G,H,I)** Z-score vector length of each recording plotted against the Pearson‘s correlation coefficient between the phase-locking strength with the mean normalised magnitude in each decile (i.e. the line in A-C except for each unit-network oscillation pair). Hollow points represent the non-significantly correlated units and filled dots represent positively correlated units. Note the positive correlation between a positive correlation and the overall phase-locking strength of the recording (Fz n=154, Pearson’s R=0.57, *P*=4.71e-15; ECoG n=109, Pearson’s R=0.55, *P*=8.67e-10; LFP n=172, Pearson’s R=0.51, *P*=6.99e-13). **(J,K,L)** Statistical comparison (MWUT) of the z-score vector length between non-correlated and significantly correlated units with network oscillation magnitude revealing a significant higher phase-locking strength of positive correlated units to all network oscillations (Fz (J), EcoG (K) and LFP (L)). This indicates that units that follow the magnitude of network oscillations have a higher phase-locking strength. *** p<0.001, box plot whiskers show 5-95th percentile.

#### Entrainment of STN-units to beta oscillations is associated with a higher oscillatory power of STN unit spike trains

Next, we investigated how the phase-locking of single neurons was related to the propensity of those neurons to oscillate at beta frequency. Higher phase-locking strength in the beta-frequency range was associated with a higher oscillatory spike train power at that frequency and neurons with significant oscillatory power had a higher phase-locking strength than non-oscillatory neurons (Supplementary Fig.1 A-C). For all field signals we observed a weak but significant correlation between degree of magnitude-dependent entrainment (R magnitude decile × z-score vector length) and oscillatory power of the spike train (Fz Pearson’s R=0.23, *P*=0.004; ECoG Pearson’s R=0.21, *P*=0.028; LFP Pearson’s R=0.39, *P*=1.20e-07; supplementary Fig. 1 D,E,F). In line with this result, the mean oscillatory spike train power in the beta frequency of preferred locking was higher for units following the instantaneous magnitude (significant for Fz and LFP), underlying the association between entrainment and oscillatory properties of the unit (comparison of spike train power in preferred beta frequency between non-correlated and positively correlated units (mean normalised magnitude decile × z-score vector length): Fz: (n=126/28), MWUT *P*=0.007; ECoG: (n=82/27), MWUT *P*=0.11; LFP: (n=116/56), MWUT *P*=2.67e-07; supplementary Fig. 1 D,E,F). Comparing the correlation strength (mean normalised magnitude at each decile × z-score vector length) between oscillatory and non-oscillatory neurons revealed a significantly higher correlation strength for oscillatory neurons for all field signals (Comparison of mean Pearson’s correlation coefficient non-oscillatory vs oscillatory units (MWUT): Fz: (n=106/48), *P*=0.007; ECoG: (n=70/39), *P*=0.002; LFP: (n=125/47), *P*=1.10e-07; supplementary Fig. 1D,E,F). These results indicate that oscillatory units show a closer relationship to the strength of ongoing beta oscillations than non-oscillatory units.

#### Subsets of STN neurons switch to a synchronised state at different beta oscillation magnitudes

Having established that phase-locking and entrainment of STN neuronal activity are positively correlated with beta oscillation magnitude, we investigated whether oscillatory neurons start phase-locking at lower magnitude values than non-oscillatory neurons. On average, significant phase-locking started around the 8^th^ magnitude decile and was lower for LFP than Fz (Mean decile at threshold of significant phase-locking: Fz 8.78 +/− 1.91 (n=74); ECoG 8.18 +/− 2.10 (n=49); LFP 7.77 +/− 2.42 (n=75); Kruskal-Wallis ANOVA F(2,192)=12.15 *P*=0.01, post hoc Dunn’s test *P*=0.01 between Fz and LFP; Fig. 5A). Non-oscillatory neurons had a higher phase-locking threshold than oscillatory units, often only showing significant phase-locking at the highest decile (Fz: non-oscillatory (n=40) 9.08 +/− 1.58, oscillatory (n=34) 8.41 +/− 2.22, MWUT *P*=0.11; ECoG: non-oscillatory (n=27) 8.85 +/− 1.51, oscillatory (n=22) 7.36 +/− 2.44, *P*=0.036; LFP: non-oscillatory (n=41) 8.85 +/− 1.59, oscillatory (n=34) 6.47 +/− 2.65, *P*=1.18e-05; Fig. 5B-D). Note that only neurons with a detectable threshold were included. The preferred frequency of phase-locking did not differ between non-oscillatory and oscillatory neurons (distribution shown in supplementary Fig. 2). While non-oscillatory STN units did not necessarily show a significant correlation with beta magnitude, the subpopulation of oscillatory units displayed a positive correlation across the entire magnitude range and their behaviour was thus highly predictable by the LFP and cortical beta activity. Therefore, subsets of STN neurons displayed different phase-locking behaviours. Non-oscillatory units tended to exhibit a discrete state change, only phase-locking to beta oscillations at the highest magnitudes or not became phase-locked at all, while oscillatory units could display a positive linear correlation across the entire magnitude range, making it possible to predict their behaviour by LFP and cortical beta activity.

**Figure 5.**
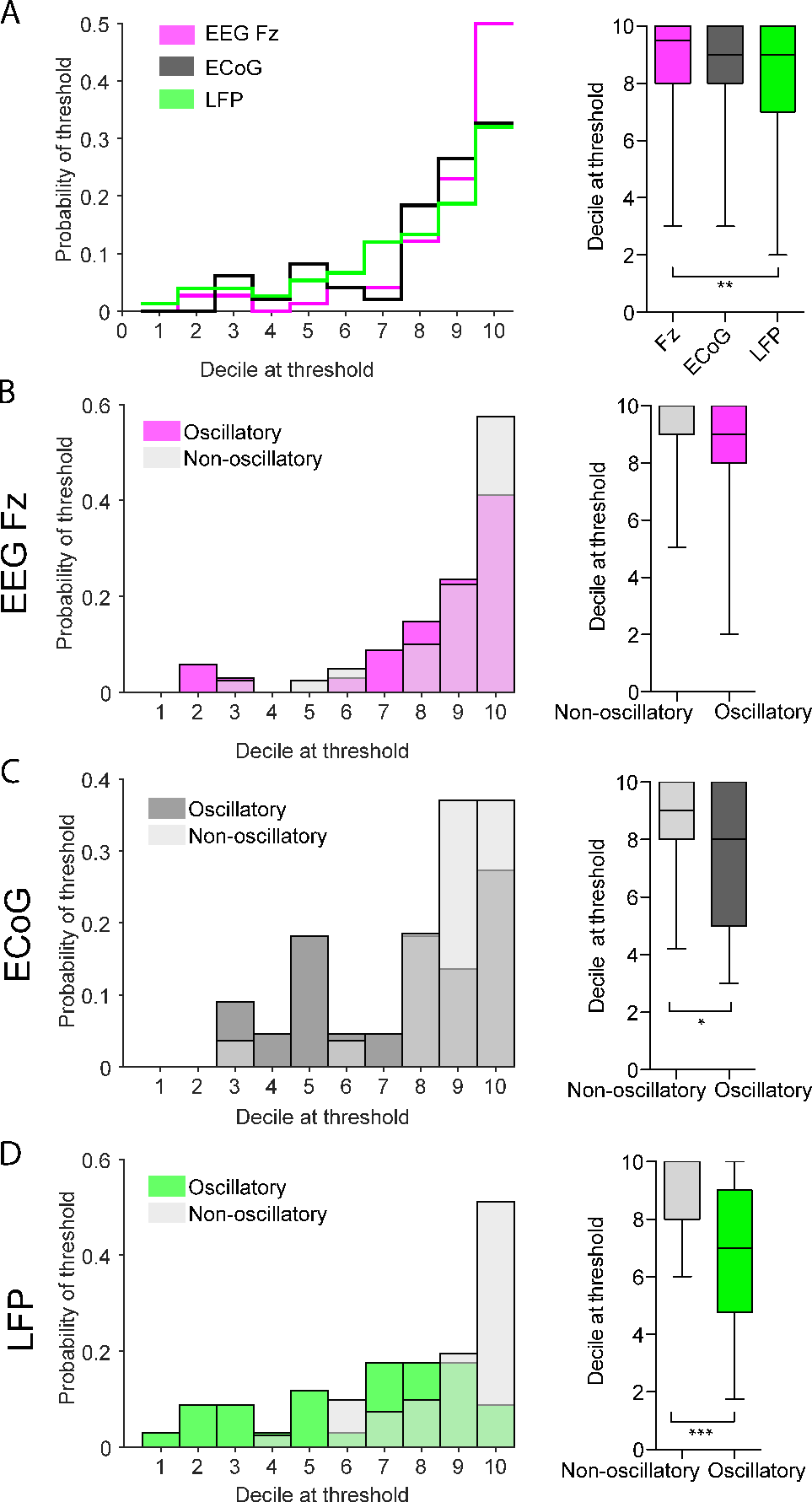
Non-oscillatory neurons require a higher magnitude of network oscillations to become significantly phase-locked. **(A)** Left, distribution of the lowest decile in which each unit becomes significantly locked to the network oscillation. Note that the likelihood increases with decile number. Right, box plots for the distributions on the left. Note the median decile of first locking is the 8th. **(B-D)** Left, distribution of lowest decile in which each unit becomes significantly locked to the network oscillations split into non-oscillatory and oscillatory neurons for Fz ((B), n=40 non-oscillatory, n=34 oscillatory), ECoG ((C), n=27 non-oscillatory, n=22 oscillatory) and LFP ((C), n=41 non-oscillatory, n=34 oscillatory). Note that the highest decile is more often the threshold or lowest decile in which significant locking starts for non-oscillatory neurons, than for oscillatory neurons. Right, boxplots of the distributions on the left. Note the decile of first significant locking is higher for non-oscillatory units for all population signals, with a significant difference for ECoG and LFP. Pairwise comparisons were performed with MWUT, *** p<0.001, *p<0.05, whiskers of boxplots show the 5-95^th^ percentile.

#### Enhanced phase-locking strength of STN-neurons during beta bursts

We have shown that the magnitude of oscillations influences the entrainment of STN-units, but next evaluated the temporal evolution of such entrainment by looking at effects during high amplitude “bursts”, which are associated with hypokinetic symptoms and can be used as a marker for therapeutic strategies (Tinkhauser G, A Pogosyan, S Little, et al. 2017; Tinkhauser G, A Pogosyan, H Tan, et al. 2017). STN units were divided in two groups, those that follow the magnitude and those that do not follow the magnitude but were significantly phase-locked. For units whose locking follows the beta magnitude, we found there was a significant difference in phase-locking before, during, and after identified bursts (Fz, F(6,336)=50.94, *P*=3.04e-09; ECoG, F(6,257)=39.38, *P*=6.02e-07; LFP, F(6,434)=53.31 *P*=1.02e-09; Kruskal-Wallis ANOVA, post hoc Dunn’s test significant values shown in Figure 6A). In contrast, the firing rate was not significantly different before, during or after identified beta burst (Fz, F(6,336)=0.28 *P*=1; ECOG, F(6,259)=0.89 *P*=0.99; LFP, F(6,441)=0.36 *P*=1).

**Figure 6.**
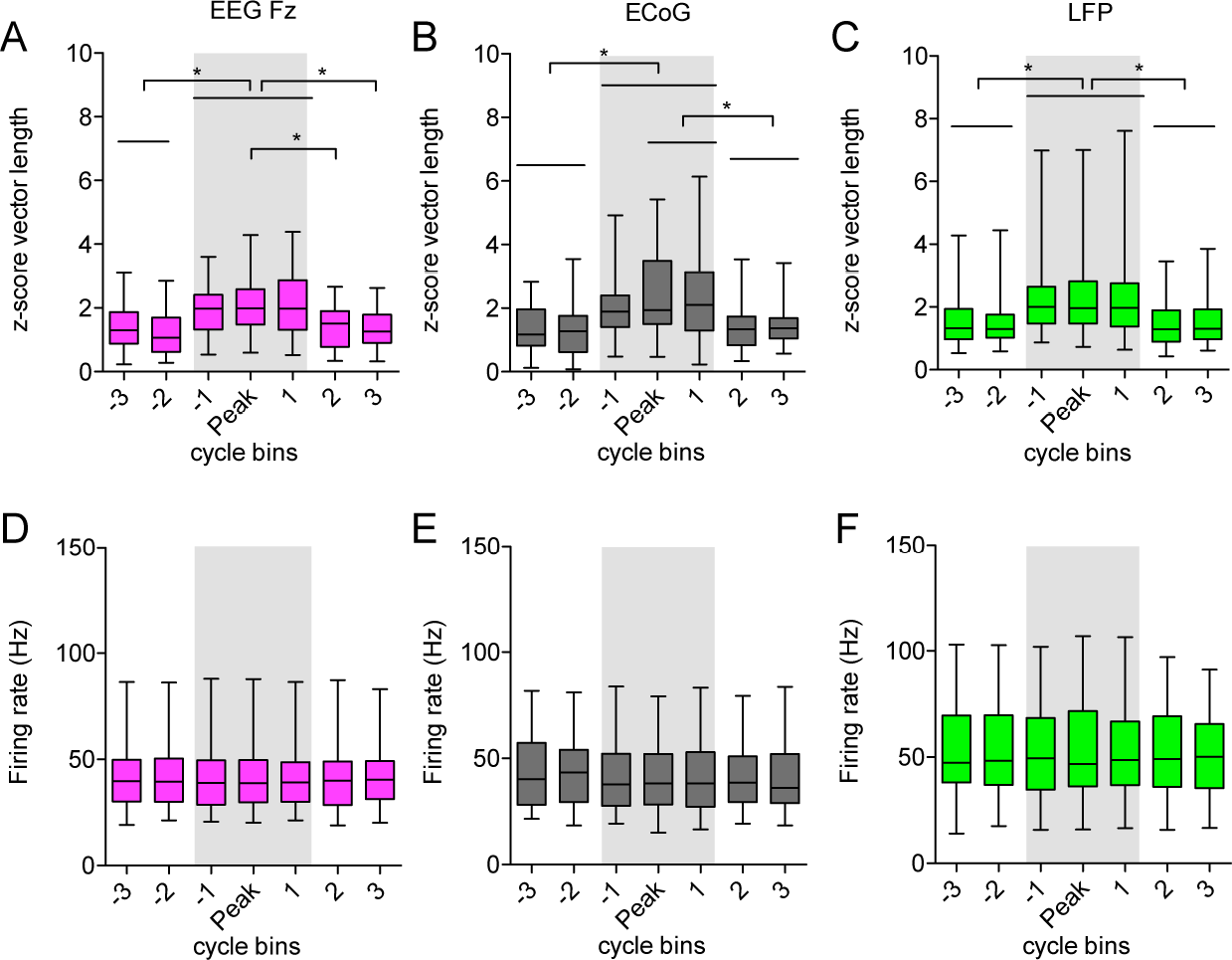
Entrainment of STN-units during periods of elevated beta power (beta bursts). **(A-C)**, Phase-locking analysis of STN-units before, during and after episodes of elevated beta power. Periods of elevated beta power were demarked using the 75th percentile theshold of the magnitude for the EEG Fz (A, pink, n=49), ECoG (B, grey,n=38) and LFP (C, green,n=64).). For each threshold crossing of more than 3 cycles the two cycles before that of the rising threshold crossing, the three cycles around the peak magnitude and the two cycles after the falling threshold crossing are depicted. Only STN units/multi-units, which showed a positive correlation of phase-locking strength with the magnitude of the oscillation are included. Note the significant higher phase-locking strength during episodes of elevated power (inside the grey shaded area). * indicate significant p-values of Dunn’s post hoc comparison. **(D-E)**, Mean firing rate in corresponding cycle bins showed in A-C for STN units during and around episodes of elevated beta power (beta bursts) in EEG Fz (D), ECoG (E) and LFP (F). A Kruskal-Wallis Anova reveals no significant difference of the firing rate within and outside a beta burst.

In contrast, STN units which did not show a significant correlation between beta magnitude and phase-locking strength showed likewise no increase of phase-locking during a beta burst (Fz F(6,666)=8.06 *P*=0.20; ECoG F(6,461)=4.90 *P*=0.56; F(6,623)=3.69 *P*=0.718) and no firing rate changes (Fz F(6,686)=0.15 *P*=0.99; ECoG F(6,476)=0.36 *P*=0.99; LFP F(6,651)=0.41 *P*=0.99) (Supplementary Fig. 3). These results indicate, that phase-locking is high during episodes of elevated beta power for a subset of significantly phase-locked STN-neurons (Fz: n=49/148 (31%); ECoG: n=38/107 (36%); LFP: n=64/158 (41%)).

### II. Entrainment of identified STN neurons to beta oscillatory input in rats

#### Cortical stimulation at different frequencies with simultaneous juxtacellular recordings in the STN of 6-OHDA hemi-lesioned rats and controls

Previous results suggest that STN neurons in PD patients are selectively sensitive to changes in the magnitude of ongoing beta oscillations in cortex. However, these analyses are dependent on natural fluctuations in the magnitude of ongoing oscillations. Moreover, we cannot determine whether this beta-selectivity is a function of the parkinsonian brain, as such data cannot be recorded from control subjects. To address these issues, experiments were carried out in anaesthetised rats, where we could manipulate the frequency of cortical input to STN neurons with a fixed magnitude in control and dopamine depleted states (Fig. 7). Approximately half of the recorded STN neurons showed a significant response (z-score>2 in the PSTH, for details see methods) in the PSTH (8/13 (62%) control, 8/17 (47%) 6-OHDA lesioned rats) to 4-40 Hz cortical stimulation (M1). Example PSTHs for a neuron from a control and a 6-OHDA lesioned rat demonstrate the type of multiphasic response described previously (Magill PJ et al. 2004) (Fig. 8). Note that typical peaks are preserved at different stimulation frequencies.

**Figure 7.**
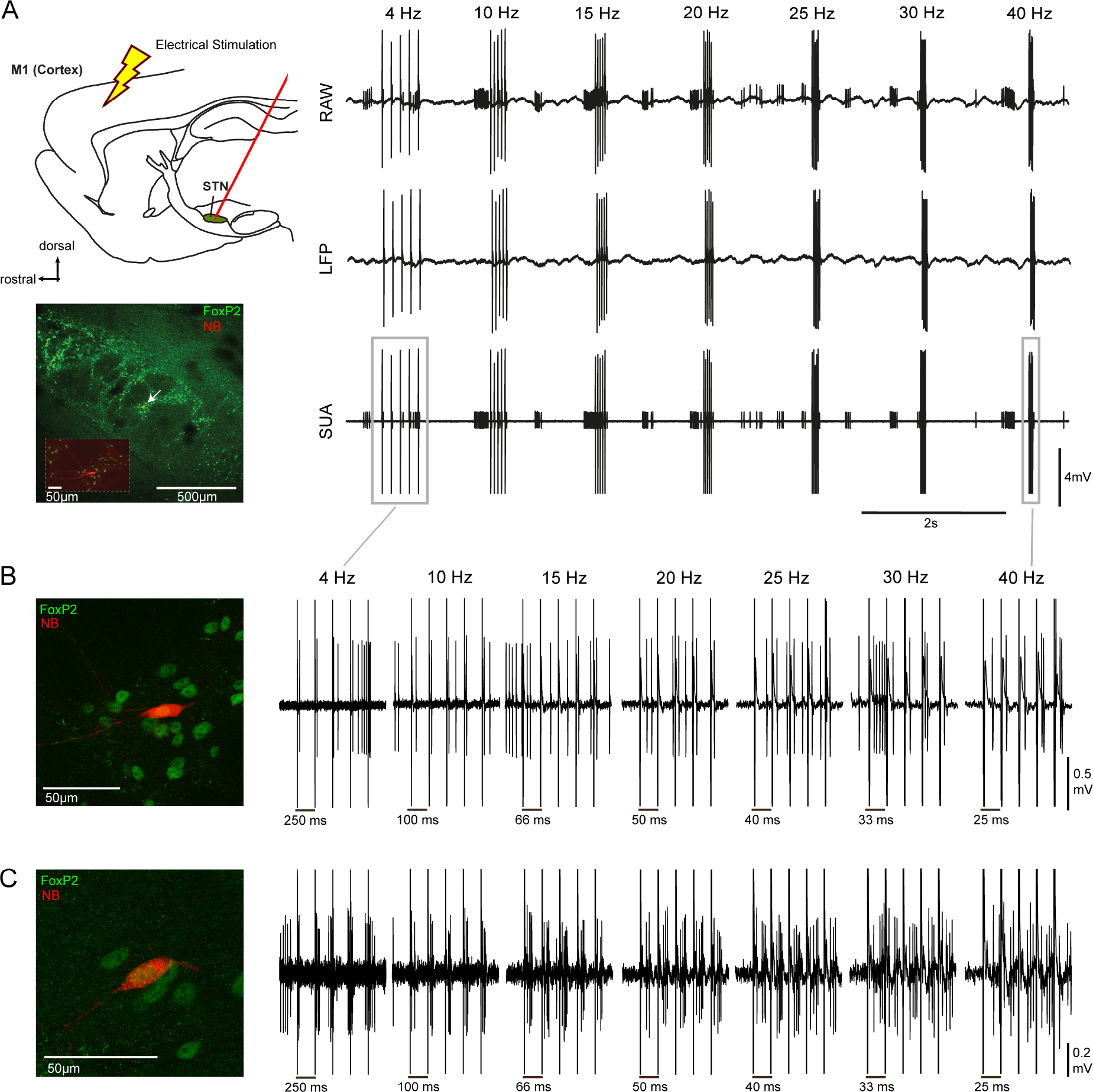
Recording setup and stimulation protocol for rat experiments. **(A)** Left top, schematic illustration of the recording setup. Recordings were performed with a glass pipette located in the STN and simultaneous electrical stimulation of the motor cortex. Left bottom, a Neurobiotin (NB) labelled neuron located in the STN, which was delineated using forkhead box protein P2 (FoxP2) immunoreactivity. A higher magnification image of the labelled neuron is inset. Right, raw recording traces from the electrode during from one block of stimulation within the 7 frequencies 4,10,15,20,25,30 and 40 Hz. For each frequency a burst of 5 consecutive stimuli was applied. This protocol was repeated 12 times. Top row, the raw signal was low-pass filtered at 2000 Hz used to extract the LFP. Middle row, the LFP signal after the removal of spikes as described in the method section. Bottom row, single unit activity was extracting following band-pass filtering between 300-5000 Hz. **(B)** The same Foxp2-expressing neuron shown in A, which was recorded in a 6-0HDA lesioned rat. Right, raw data recorded from the labelled neuron. The period around each burst of stimuli for each individual frequency has been magnified. Note the time scale of the plots differs. In this example, 4 Hz stimulation results in the timing the evoked spikes having high variance over the course of the stimuli, which is reduced at higher frequencies **(C)** Left, a FoxP2-expressing labelled neuron from a healthy rat. Right, the corresponding raw data traces from the labelled neuron. Note the similar entrainment pattern over course of stimuli as shown in B.

#### STN-neurons in the cortico-pallidal-subthalamic network show frequency selective entrainment to cortical stimulation in healthy and parkinsonian rats

To measure neuronal synchronisation to a given stimulus frequency, we calculated the Coefficient of Variation (CV) of the first spike occurring after each stimulus pulse. Rasterplots of example neurons show the increasing consistency in the timing of the first evoked spike at 10, 15, 20 and 25 Hz (Fig. 8), suggesting increasing synchronisation over consecutive pulses. Because we could not observe obvious differences in the single unit responses (PSTHs) between 6-OHDA lesioned (n=8 STN neurons) and control (n=8 STN neurons) animals, groups were combined to increase statistical power (n=16 STN neurons).

**Figure 8.**
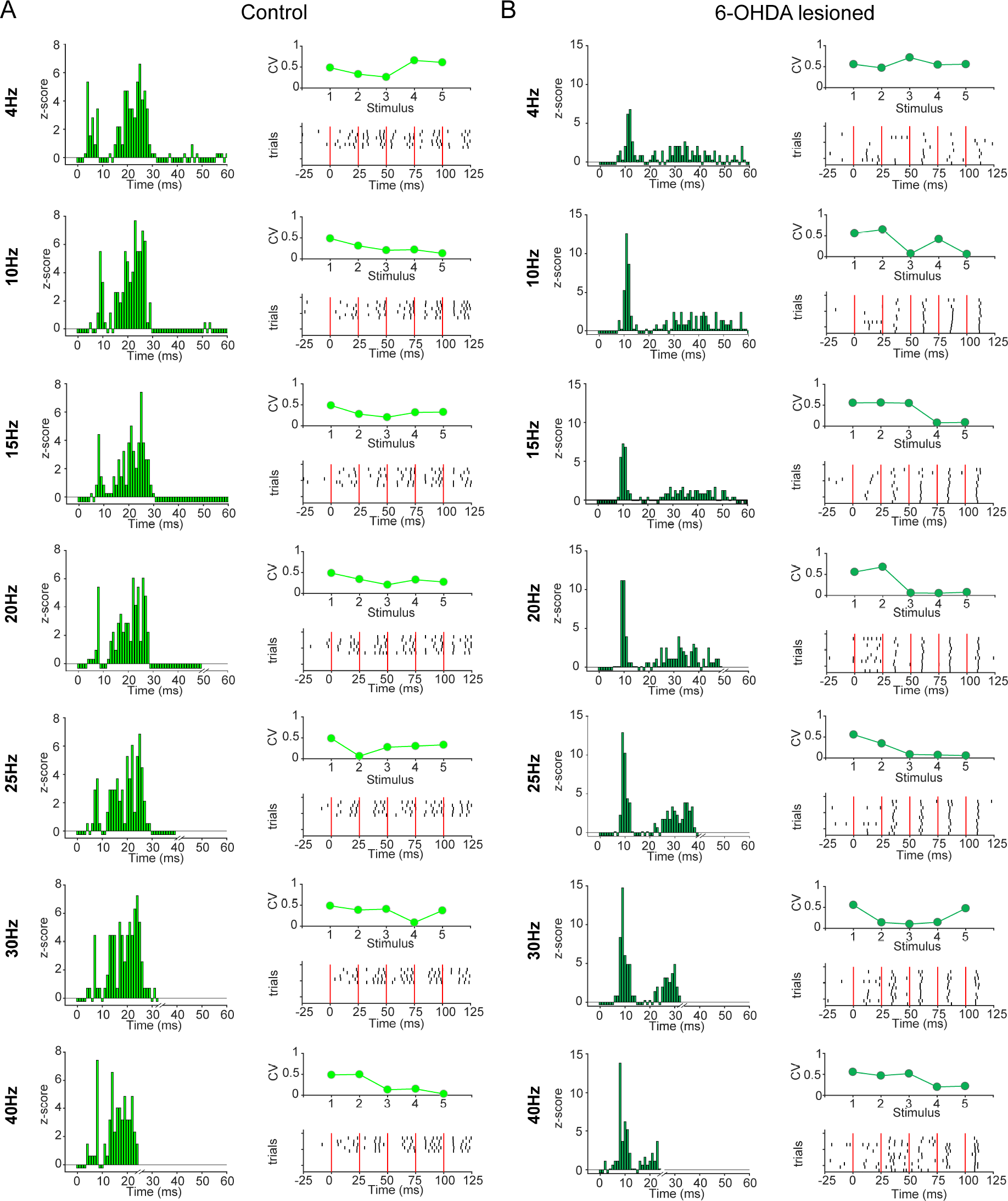
Example responses of identified STN-neurons to cortical stimulation at different frequencies. Each column shows the response of an STN neuron from a control rat **(A)** and 6-OHDA hemi-lesioned rat **(B)** to alternating stimulation in the frequencies 4, 10, 15, 20, 25, 30 and 40 Hz (rows) with 5 consecutive stimuli during one recording session. Left, Normalised PSTH (z-score, based on 500 ms of pre-stimulus period) to the stimulation at time 0. All consecutive stimuli were merged for the calculation of the PSTH. Note that the observation window decreases with the stimulation cycle period. Right bottom, Rasterplot of all available trials for the STN neuron. The green line indicates the stimulus timing, with in total 5 consecutive stimuli per trial. Each line represents 1 trial in the recording session. Note that the time scale was matched to be equivalent for all frequencies and therefore for lower frequencies not the whole post-stimulus period is shown. Right Top, Coefficient of Variation (CV) across trials of the first spike after the stimulus indicating the amount of synchronisation to the stimulus. Note the decreasing Coefficient of Variation (CV) in the frequencies 10-25 Hz in both examples, indicating an increasing synchronisation to the stimulus with increasing stimulation duration.

Across all neurons the CV decreased with the course of consecutive stimuli selectively in response to beta-frequency (10-25 Hz) cortical stimulation (4 Hz: F(4,60)=7.25, *P*=0.12; 10 Hz: F(4,60)=19.6, *P*=0.0006; 15 Hz: F(4,60)=12.44, *P*=0.0005; 20 Hz: F(4,60)=14.65, *P*=0.006; 25 Hz: F(4,60)=11.45, *P*=0.022; 30 Hz: F(4,60)=2.35, *P*=0.67; 40 Hz: F(4,44)=7.2, *P*=0.13; Friedman test, significant post hoc comparisons are stated in Fig. 9 and in supplementary Table 2). These results suggest STN unit entrainment becomes stronger over the duration of the stimulus train in the frequencies 10-25 Hz. In support of this conclusion, significant decreases in CV following the 1^st^ vs 5^th^ stimulus within each frequency occurred for 10, 15, 20 and 25 Hz, but not other frequencies (Signed Wilcoxon rank test: n=16 STN neurons; 4 Hz: *P*=0.35; 10 Hz: *P*=9.35e-04; 15 Hz: *P*=0.001; 20 Hz *P*=0.011; 25 Hz *P*=0.020; 30 Hz *P*=0.76; 40 Hz *P*=0.54).

**Figure 9.**
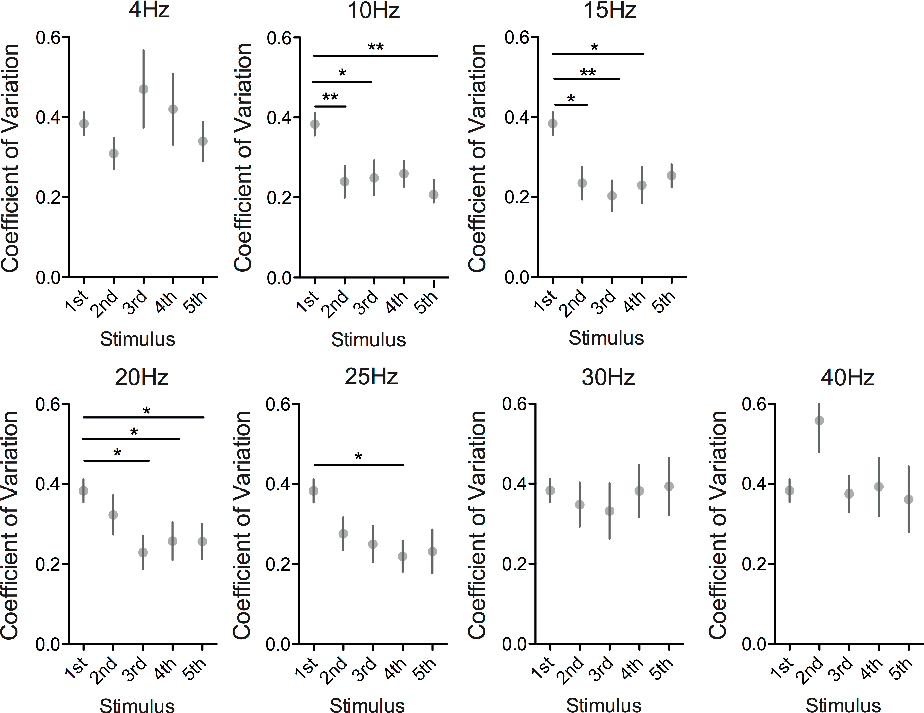
Frequency selective entrainment of identified STN-neurons to cortical stimulation. Group analysis for the Coefficient of Variation (CV) of the first spike after each stimulus across trials. In this sense the CV is a measurement of spike synchronisation to the stimulus. All spike times with respect to all available first stimulus across frequencies have been used to calculate the CV of the first stimulus because the stimulation frequency is defined only with the second pulse. Each subplot shows the results across stimuli for the stimulation frequencies 4 Hz, 10 Hz, 15 Hz, 20 Hz, 25 Hz, 30 Hz and 40 Hz. Dots show group mean +/− SEM. Stars indicate p-values of Tukey‘s post hoc comparison after a Friedman test. *p<0.05, **p<0.01. Note the significant difference for 10 and 15 Hz already after the second pulse, indicating an immediate higher degree of synchronisation at those frequencies. In addition, the entrainment at 20 and 25 Hz seems to follow more a gradient pattern. 4, 30 and 40 Hz showed no improvement of synchronisation across the course of stimuli.

When control and lesion groups were separated, results were similar (Supplemental Fig. 4). As above, a significant decrease in CV following the 1^st^ and 5^th^ stimulus occurred for 10 and 15 Hz in both controls (n=8, 10 Hz: *P*=0.008, 15 Hz: *P*=0.039) and 6-OHDA lesioned animals (n=8, 10 Hz: *P*=0.039, 15 Hz: *P*=0.016). In addition, 6-OHDA lesioned animals showed a significant difference at 25 Hz (n=8, *P*=0.023), although there was a similar trend in controls (Supplementary Fig. 4). Note that even with split groups there was a tendency of the 5^th^ stimulus CV to be lower for the frequencies 10-25 Hz (Supplementary Fig. 4). The tendency of STN neurons to be entrained by beta-frequency cortical input was therefore not a specific property of dopamine depleted animals.

#### A decrease in excitability accompanies increased synchronisation to cortical stimulation at beta frequencies

Having shown that STN neurons are more precisely timed by beta-frequency input, we aimed to elucidate whether this was associated with changes in firing rate. Figure 10A shows the firing probability across stimuli for each frequency. At 4Hz and 10Hz, stimulation led to a significant increase in firing rate (4Hz: F(4,60)=16.47 *P*=0.003; 10 Hz: F(4,60)=10.45 *P*=0.034), and for 15 Hz (F(4,60)=6.32 *P*=0.18) and 40 Hz (F(4,60)=6.68 *P*=0.15) no significant difference was observed. In contrast, the 20-30Hz stimulation showed a decrease in firing probability across the stimulus train (20Hz: F(4,60)=26.37 *P*=0.003; 25 Hz: F(4,60)=22.95 *P*=0.0001; 30 Hz: F(4,60)=16.69 *P*=0.002; significant results of post hoc comparisons are stated in Fig. 10 and supplementary Table 3). Interestingly, in addition to this decrease in firing probability, there was an increase in spike latency across the stimulus train at frequencies from 10 to 30Hz (Supplementary Fig. 5 and supplementary Table 4).

**Figure 10.**
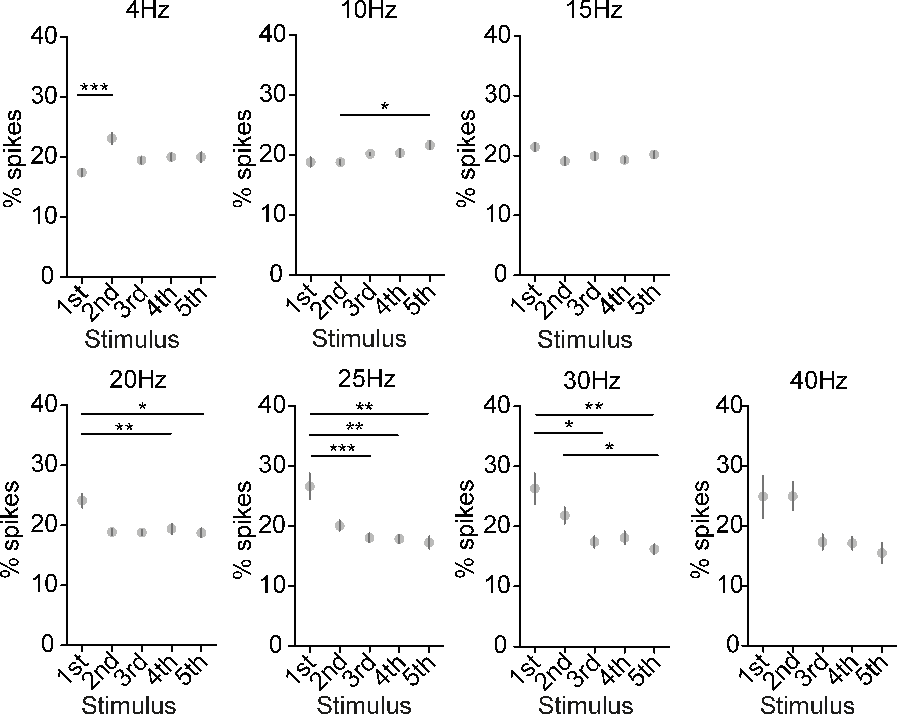
Decrease of firing probability of STN neurons over the course of cortical stimulation at beta frequencies. Firing probability after each stimulus expressed in % spikes after each stimulus with respect to all spikes during the burst of stimuli. Each subplot shows the firing probability over the course of stimuli at one frequency (at 4, 10, 15, 20, 25 and 30 Hz). Dots show the mean +/− SEM of the group (n=16 STN neurons). Note the significant decrease of firing probability already after the 2nd pulse for 20 Hz and the more gradually decrease in the frequencies 25 and 30 Hz. This indicates that selectively in the beta-frequency range the firing probability gets suppressed. Overall, these results suggest an inhibition of spiking at the frequencies 10-30 Hz, which might even increase with additive stimuli. For comparison, a Friedman test with post hoc Tukey’s comparisons was performed. *p<0.05, **p<0.01 and ***p<0.001

Because the difference between the 5^th^ and 1^st^ stimulus appeared to be highest, we subtracted the value of the 1^st^ stimulus from the 5^th^ stimulus for the parameters CV, % spikes and spike latency after the stimulus and compared it across frequencies. The CV-difference appeared to be lower in the frequencies 10-25 Hz but was only significantly different from 30 Hz for 10-20 Hz (Supplementary Fig. 6A, F(6, 78)=15.64, *P*=0.02, post hoc Tukey’s 10-30Hz *P*=0.02, 15-30Hz *P*=0.046, 20-30Hz *P*=0.046). Spike suppression over the course of stimuli was more likely in the frequencies 15-30Hz in comparison to 4 and 10Hz (Supplementary Fig. 6B, F(6,90)=40.24, *P*=4.08e-07, post hoc Tukey’s 4Hz-20Hz *P*=0.025, 4Hz–25Hz *P*=0.003, 4Hz-30Hz *P*=0.001, 10Hz-25Hz *P*=0.010, 10Hz-30Hz *P*=0.005). This indicates a suppression of spike probability over the course stimuli in the frequencies 10-30 Hz in comparison to low frequency stimulation. This was supported by the tendency of an elevated difference in spike latency in the frequencies 10-30 Hz. The result of 25 and 30Hz stand out with a significant difference between 4 Hz and 25/30 Hz (F(6,90)=21.21, *P*=0.002, post hoc Tukey’s 4Hz-25Hz *P*=0.006, 4Hz-30Hz *P*=0.03). In summary, these changes suggest that beta-frequency input supresses and synchronises the spiking of a STN-neuron in contrast to 4 or 40 Hz stimulation.

#### Altered STN evoked responses in the 6-OHDA lesioned rat suggest decreased pallidal reciprocal inhibition

The timing and pattern of peaks in the PSTH and evoked LFP following cortical stimulation give an estimate of different pathways through which the cortical excitation reaches the STN (Magill PJ et al. 2004). Peaks in the PSTH are reflected in corresponding troughs in the LFP, while the first peak in the evoked LFP response (P1) reflects early inhibition by reciprocal GPe connections (Magill PJ et al. 2004). Figure 11 shows the averaged PSTH for each frequency with two prominent peaks corresponding to the early and late excitation in both groups. Interestingly, on average the peak is delayed in 6-OHDA lesioned animals, in line with a down-regulation of the hyper-direct pathway (Chu HY et al. 2017; Wang YY et al. 2018). Moreover, the mean z-score PSTH does not decrease to 0 in the 6-OHDA lesioned animals, an effect that was magnified as input frequency increased, supporting the idea of a decreased reciprocal inhibition from the GPe in the dopamine depleted state (Janssen MLF et al. 2017).

In accordance with this result, averaged LFP evoked responses in the 6-OHDA lesioned animals show delayed N2, P1, and P2 peaks (Fig. 10). Evoked responses in STN unit firing can express a variety of patterns (Kita H and T Kita 2011). In our data 6/12 evoked LFP responses had a flat response, while this happened only in 2/12 neurons in control animals. All other individual evoked responses had at least a clearly identifiable P1. Because peak height difference between control and 6-OHDA lesioned was most pronounced in the time window 5-15 ms, we calculated the mean z-score LFP amplitude in that window for each recording and compared it between the groups (Fig. 11) (MWUT, n=12 recordings per group, 4Hz: *P*=0.708; 10Hz: *P*=0.750; 15Hz: *P*=0.341; 20 Hz: *P*=0.035; 25 Hz: *P*=0.009; 30 Hz: *P*=0.002; 40 Hz: *P*=7.10e-04). Notably, the difference was only significantly different in the frequencies 20, 25, 30 and 40 Hz. These changes provide further evidence that STN-driven reciprocal GPe inhibition is reduced in the dopamine depleted brain. These changes, however, do not seem to completely underlie the locking of single neurons to beta-frequency input, which was seen in both healthy and parkinsonian animals.

**Figure 11.**
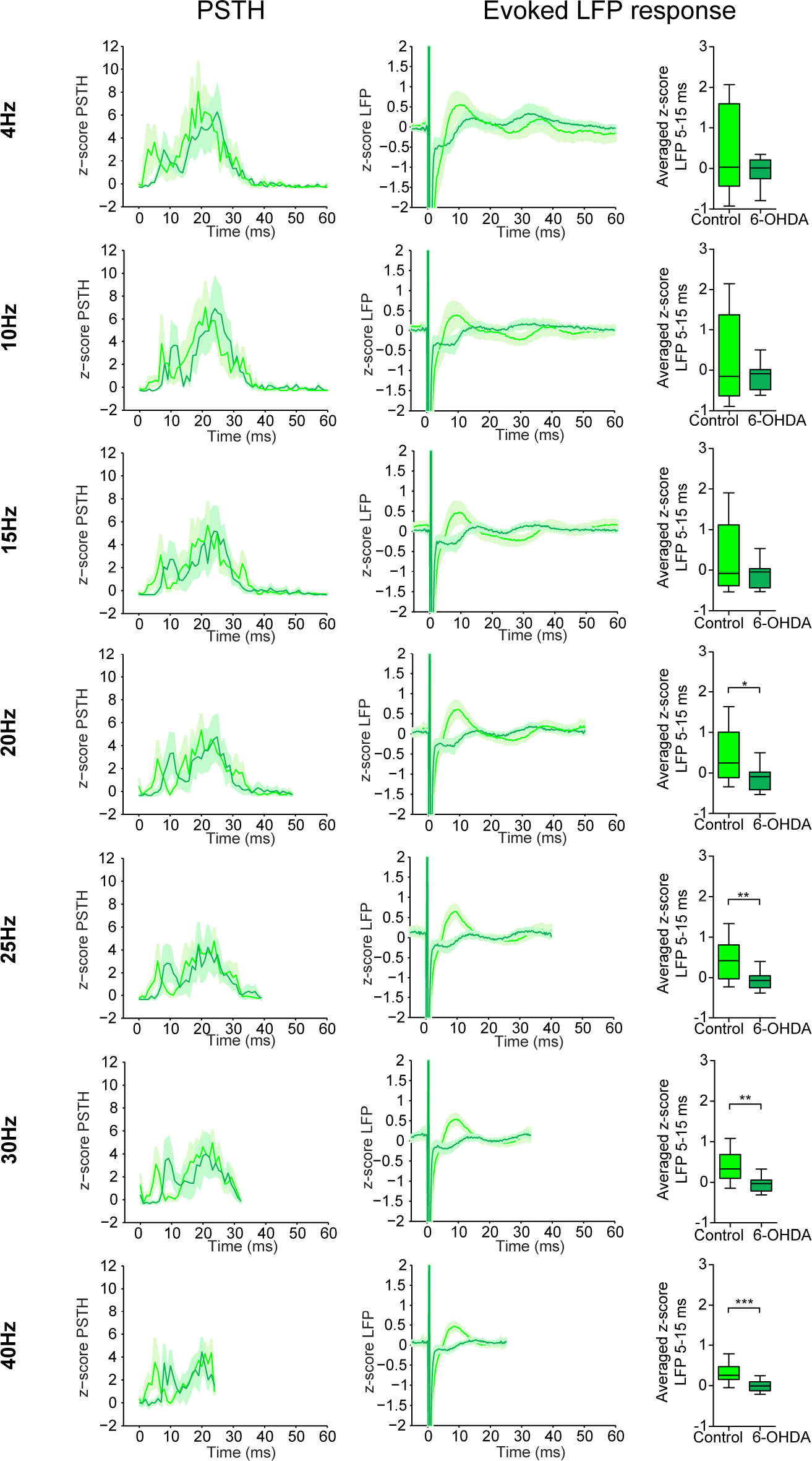
Comparison of the evoked LFP-response between control and 6-OHDA lesioned rats. Left, mean z-score PSTH (PSTH normalised for the 500 ms pre-stimulus period) with SEM shading for all stimuli in each frequency for STN-neurons expressing a significant response in the PSTH in control (n=8, light grey) and 6-OHDA (n=8, dark grey) lesioned animals. Each row represents 1 frequency stated on the left. Middle, Mean evoked LFP-response of STN-LFP (z-score was calculated by subtracting the mean and dividing it by the standard deviation of the mean amplitude for each recording) to cortical stimulation. For the LFP analysis we used stimulation amplitude matched unique recording positions for control (n=12, light grey) and 6-OHDA lesioned (n=12, dark grey) rats. Note the similar shape of the mean curve for each group across frequencies with a more pronounced effect in the frequencies 20-30 Hz. Moreover, note the time shift in accordance of the PSTH peaks with a peak delay in 6-OHDA rats for all peaks. But in addition, even with the time delay of the P1, the group peak in the 6-OHDA seems to be less pronounced in comparison to controls, indicating a possible reduced reciprocal inhibition from the GPe. Right, boxplots show a comparison of the mean z-score of the LFP amplitude in the time interval 5-15 ms for each frequency (stated on the left side of the row). Note the significant difference in amplitude for 25 and 30 Hz in MWUT. *p<0.05, **p<0.01,***p<0.001. Boxplots show 5-95 percentile.

## Discussion

The phase-locking of neuronal activity in the STN, and other basal ganglia populations, to beta-frequency network oscillations in PD is thought to limit information coding space in these circuits, ultimately leading to akinetic/rigid symptoms (Engel AK and P Fries 2010; Brittain JS et al. 2014; Khanna P and JM Carmena 2015). It remains unclear, however, why these networks resonate at beta frequency and whether this frequency selectivity is engrained into the system, or is purely pathological. We address this question by combining STN spiking data from PD patients with experiments where it was possible to manipulate the frequency of cortical input to STN in experimental animals. Together, our results suggest that STN neurons are preferentially driven by beta-frequency input in PD, and that this preference may be due to the connectivity of the microcircuit in which they are embedded.

### Frequency-selective and magnitude-dependent entrainment of STN neurons to network oscillations in PD patients

Synchronisation of STN neurons to local and cortical beta oscillations in PD patients has been extensively described (Kuhn AA et al. 2005; Weinberger M et al. 2006; Moran A et al. 2008; Sharott A et al. 2018). Phase-locking provides an important metric, as it indicates that individual neurons have become engaged with activities that are synchronised across the network. Indeed, the extent and magnitude of phase-locking between STN units and frontal cortical signals correlates with motor impairment (Sharott A et al. 2018). While STN neurons lock to frequencies across a wide beta range (Shimamoto SA et al. 2013; Sharott A et al. 2018), we found that they rarely do so with theta/alpha and gamma oscillations. The relatively low incidence of locking to theta/alpha oscillations is surprising, given the higher or equivalent power of these activities in the cortex and STN LFP. The selectivity of STN units to beta frequencies does not, therefore, appear to be a trivial result of their increased power in PD and likely reflects some property of STN neurons or the network in which they are embedded.

It has been previously shown that the magnitude of network oscillations in the motor cortex drives the engagement of cortical neurons with LFP beta oscillations (Denker M et al. 2008). We found that phase-locked STN neurons displayed heterogeneous relationships with ongoing amplitude of network oscillations. Some neurons had a positive linear relationship, or transfer function, between instantaneous beta amplitude and phase-locking. Importantly, these neurons were most likely to oscillate at beta-frequency, and thus be of particular relevance to akinetic/rigid disease symptoms (Sharott et al 2014). The full dynamic range of LFP amplitude, therefore, gives the most information about the neurons that are most strongly related to disease pathology. Other neurons had a weaker relationship to instantaneous amplitude, significantly phase-locking only at the highest amplitudes. Periods of high beta-amplitude thus represent epochs during which an additional group of neurons are also recruited in to the network oscillation. We saw little evidence of phase-locking of STN neurons to other frequencies, even at these highest amplitudes, indicating that beta selectivity is maintained even at maximum input. Finally, some phase-locked neurons had no observable relationship with instantaneous amplitude. Here, we cannot rule out that we were possibly recording a suboptimal population signal for those particular units, for example an unconnected cortical area or too distant LFP recording position. However, as the spiking of these neurons was phase-locked to that signal over the entire recording, this result suggests that some neurons are only very weakly coupled to the network oscillation.

### Temporal dynamics of neuronal synchronisation in the STN

The instantaneous amplitude of beta oscillations has gained increasing importance given the recent focus on the transient nature of beta oscillations in sensorimotor circuits, often referred to as “beta bursts,” in both health (Feingold J et al. 2015; Sherman MA et al. 2016; Khanna P and JM Carmena 2017; Shin H et al. 2017) and disease conditions (Tinkhauser G, A Pogosyan, S Little, et al. 2017; Tinkhauser G, A Pogosyan, H Tan, et al. 2017; Lofredi R et al. 2018; Tinkhauser G et al. 2018; Torrecillos F et al. 2018). In PD, metrics of beta bursts are correlated with clinical parameters and have provided an effective biomarker for “adaptive” DBS, whereby high frequency DBS is triggered by the presence of a burst or high amplitude (Little S et al. 2013; Arlotti M et al. 2018). While these studies have provided compelling evidence that beta bursts can predict motor and disease-related behaviour, they have generally not demonstrated how such events influence underlying processing by individual neurons. In particular, it is not clear whether thresholding the ongoing power represents a point in a continuum or a step change in the engagement of the underlying neurons with an incoming oscillatory input and/or local synchronisation. This relationship between the LFP and spiking is crucial, as it will ultimately dictate the impact of the oscillation on coding within STN and in downstream structures.

Our analysis of the relationship between LFP/EEG amplitude and phase-locking suggests that, in the STN, commonly used thresholds appear well designed to capture maximum engagement units to population oscillations. Here we provide the first evidence in humans that STN LFP and cortical beta bursts predict periods of phase-locking in single and multiunit activities. Our analyses also show, however, that such thresholds may exclude valuable information that occurs at lower amplitudes. This finding has important implications for the design of closed-loop stimulation for the treatment of PD. A high threshold may well be optimal for adaptive DBS, where the aim is to disrupt periods of maximum synchronisation as they occur. For other approaches, it could be possible to utilise the lower amplitude information. Several authors have postulated that phase-dependent stimulation could be an effective therapeutic strategy for treating motor symptoms (Rosin B et al. 2011; Azodi-Avval R and A Gharabaghi 2015; Holt AB et al. 2016; Meidahl AC et al. 2017) and we have recently demonstrated the potential efficacy of this approach (Holt AB et al. 2018). Together, these studies show that it may be possible to utilise phase-dependent stimulation at low amplitudes, disrupting the most synchronised population of neurons before pathologically sustained beta bursts arise.

### Beta frequency cortical input selectively reduces spike-time variance

Our analysis of ongoing oscillatory activity suggests that STN neurons are selectively responsive to beta-frequency input. However, it is impossible to fully test this using spontaneous data where certain frequencies, such gamma, do not reach equivalent power to those in the lower bands. Using cortical stimulation in rodents allowed us to provide a controlled oscillatory input of equal amplitude, but variable frequency, to identified STN neurons in both healthy and dopamine-depleted states. While such input is undoubtedly far stronger than naturally occurring beta bursts, such experiments allowed us to dissect the response of STN neurons with high temporal precision. Input at beta, but not other frequencies, decreased the variability of the timing of evoked spikes, consistent with a higher fidelity of locking of the STN neurons to that input in patient data.

Given that we observed a concurrent reduction in the number of spikes fired and an increase in spike latency during beta stimulation, a trivial explanation for our findings is that intrinsic properties of STN neurons simply cannot follow beta frequencies, reducing variance in spike timing. Several lines of evidence suggest that this is not the case. Firstly, the highest frequency used here (40Hz) did not lead to any delay in spike initiation between the first and later stimulus pulses. Secondly, EPSPs in STN neurons reduce, rather than increase, the firing threshold immediately after a spike is fired (Farries MA et al. 2010). Finally, STN neurons can follow far higher frequency inputs (Bevan MD and CJ Wilson 1999; Do MT and BP Bean 2003), making it unlikely that their intrinsic ability to follow the frequencies used here is reduced over successive stimuli. Indeed, STN neurons do not display any specific resonance in their intrinsic properties (Farries MA et al. 2010).

While the in vivo recordings used here do not allow us to examine these intrinsic dynamics directly, they have the advantage allowing the measurement of the STN neuron response in the intact network. Moreover, the multiphasic response of STN, and other basal ganglia nuclei to single pulses of cortical stimulation has been extensively described (Nambu A et al. 2000; Magill PJ et al. 2004; Farries MA et al. 2010; Janssen MLF et al. 2017) and there is an established framework for the mechanisms thought to underlie each phase (Magill PJ et al. 2004). In particular, the first three phases of the response correspond to hyperdirect input, which drives GPe mediated inhibition and is followed by a rebound excitation (Nambu A et al. 2000; Farries MA et al. 2010). Beta-frequency input (40-66ms interval) would arrive just after this hyperdirect-driven polysynaptic response and approximately at the time when indirect pathway-driven inhibition arrives (Nambu A et al. 2000; Magill PJ et al. 2004; Farries MA et al. 2010; Nishibayashi H et al. 2011).

In STN neurons, the simultaneous arrival of IPSPs is necessary for oscillatory EPSPs to entrain spiking at the input frequency (Baufreton J et al. 2005). Bevan and colleagues proposed that the simultaneous IPSPs would result from the STN of the reciprocal pathway with GPe (Baufreton J et al. 2005; Baufreton J et al. 2009). Here we show that in dopamine-lesioned animals, this reciprocal inhibition is greatly reduced, as originally suggested by Benazzouz and colleagues (Janssen MLF et al. 2017). It seems unlikely, therefore, that this pathway is necessary for the beta-selectivity seen during cortical stimulation, as the reduction of this reciprocal inhibition in lesion animals did not reduce the entrainment by cortical input as compared to controls. In addition, recent findings show that that the hyperdirect pathway, and presumably its ability to drive GPe, is weakened in PD (Chu HY et al. 2017).

We hypothesise that a coincident inhibitory drive *is* necessary to entrain STN neurons to cortical input in vivo, but is mediated by the indirect, rather than the hyperdirect pathway. For spontaneous, cortically driven oscillations in PD, the hyperexcitability of indirect pathway spiny projection neurons would allow this simultaneous entrainment of GP by cortical input, which would usually be dampened in the presence of dopamine (Cagnan H et al. 2018; Sharott A et al. 2018). In healthy animals, STN neurons display little or no oscillation at beta frequency (Mallet N, A Pogosyan, A Sharott, et al. 2008). Using cortical stimulation, however, beta-frequency input could also preferentially entrain STN in healthy animals. This could be because cortical stimulation artificially drives striatal projection neurons, even in the presence of dopamine (Mallet N et al. 2005; Sharott A et al. 2012), creating a network drive more similar to that in PD. As selective entrainment at beta frequency still can still occur under control conditions it could at least partly be a function of the dynamics of the healthy connectivity, rather than a result of pathological plasticity. The relative quiescence of the healthy striatum could act to maintain decorrelation in the STN (Wilson CJ 2013), by preventing indirect pathway input synchronising with that of the hyperdirect pathway. If this is the case, novel therapeutic strategies that prevent rhythmic indirect pathway input to STN could tip the balance back towards intrinsic firing and reduce motor impairment.

Overall, our findings are in line with recent work by Bevan and colleagues, demonstrating that the key pathological feature of the STN in PD could be the change in the ratio of intrinsically driven to input driven spiking (McIver EL et al. 2018). Beta-frequency synchronisation between cortex and STN could be the most common manifestation of the disruption of this ratio due to an underlying selectivity for these frequencies in the surrounding microcircuit.

## Acknowledgements

The authors would like to thank all the patients who participated in this study.

## Supplementary Material

**Supplementary Figure 1.**
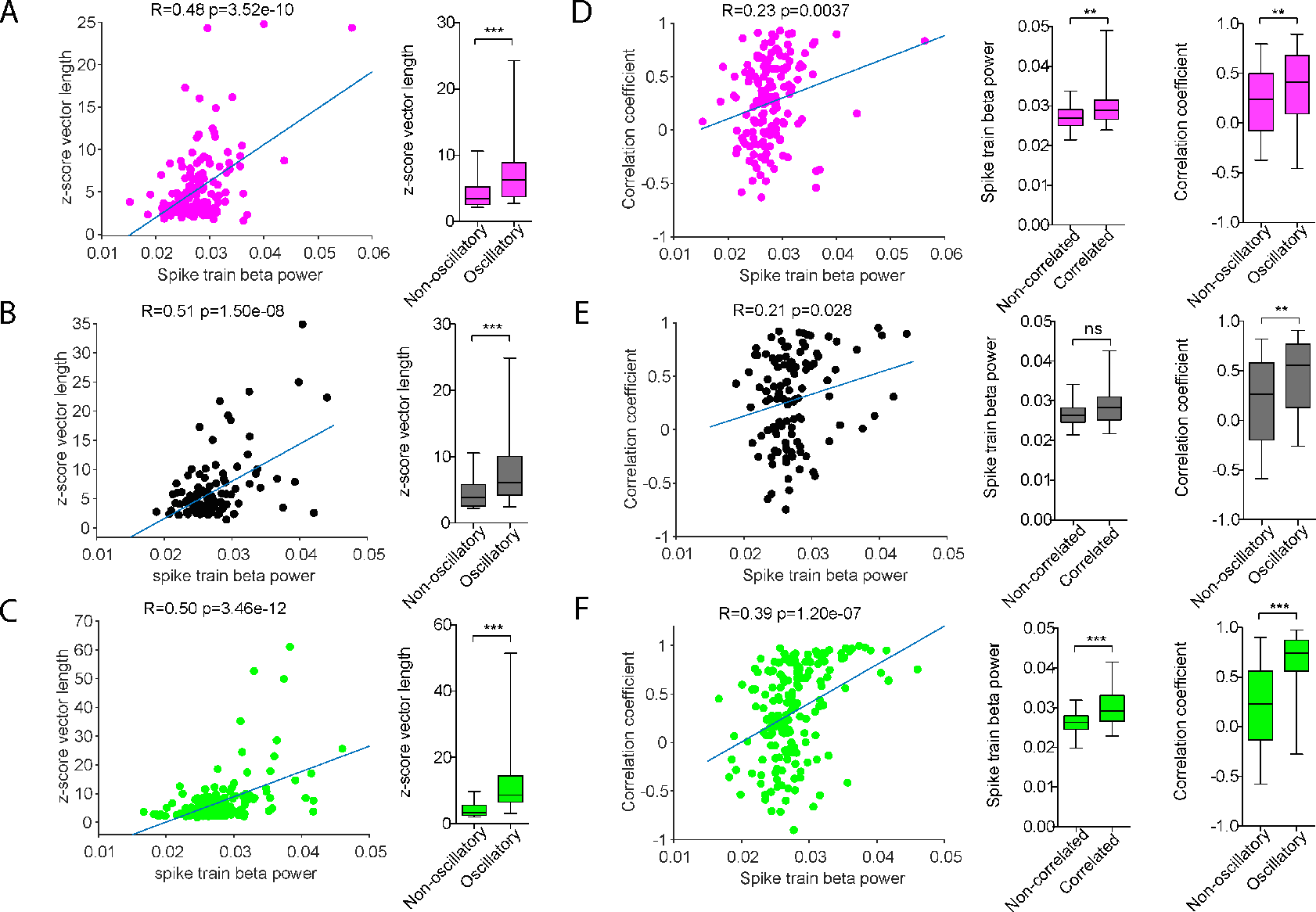
Phase-locking and entrainment of STN units to beta oscillations is associated with a higher oscillation strength of STN units. **(A-C)** Dots in the scatter plot show the mean z-score vector length (phase-locking strength) at the beta frequency of preferred phase-locking (12-40 Hz) and spike train beta power at the same frequency +/− 5 Hz normalised by the spike train power in the range 300-500 Hz for each recorded pair. The phase-locking strength is significantly positively correlated with the normalized spike train power for all investigated signals (Fz Pearson’s R=0.48, *P*=3.52e-10; ECoG Pearson’s R=0.51, *P*=1.50e-08; LFP Pearson’s R=0.50, *P*=3.46e-12). The line indicates the linear fit of the correlation. The boxplots on the right hand side show the z-score vector length for non-oscillatory and oscillatory neurons with a significant higher phase-locking strength for oscillatory neurons for all investigated network oscillation signals (Fz (A), ECoG (B) and LFP (C). (Fz: non-oscillatory units (n=106/154), oscillatory units (n=48/154), *P*=1.82e-06; ECoG: non-oscillatory units (n=70/109), oscillatory units (n=39/109), *P=8*.32e-05; LFP: non-oscillatory units (n=125/172), oscillatory units (n=47/172), *P=1.39*e-12)). **(D-F)** Show the relationship between entrainment of STN unit to beta oscillations (measured by the Pearson’s correlation coefficient between the phase-locking strength and the mean magnitude at each percentile) and oscillatory properties of the STN units for all network oscillation signals EEG Fz (D), ECoG (E) and LFP (F). Left, Scatterplots show the Pearson’s correlation between the Pearson’s correlation coefficient (z-score vector length × mean normalised magnitude at each percentile) and the spike train power in the preferred frequency of phase-locking +/− 5 Hz analogue to A-C. Note the significant positive correlation between magnitude-dependent phase-locking and oscillation strength for pairs with all network oscillation signals (Fz Pearson’s R=0.23, *P=0.004*; ECoG Pearson’s R=0.21, *P*=0.028; LFP Pearson’s R=0.39, *P*=1.20e-07). Middle, Boxplots show that the normalized spike train beta power (as described above) is higher for significantly positive correlated pairs (Fz: non-correlated units (n=126/154), correlated units (n=28/154), *P*=0.007; ECoG: non-correlated units (n=82/109), correlated units (n=27/109), *P=0.11*; LFP: non-correlated units (n=116/172), correlated units (n=56/172), *P=2.67*e-07). Right, Boxplots show the Pearson’s correlation coefficient (z-score vector length × mean normalised magnitude at percentile) for non-oscillatory and oscillatory neurons (Fz: non-oscillatory units (n=106/154), oscillatory units (n=48/154), *P*=0.007; ECoG: non-oscillatory units (n=70/109), oscillatory units (n=39/109), *P=0.002*; LFP: non-oscillatory units (n=125/172), oscillatory units (n=47/172), *P=1.10*e-07). This indicates, that oscillatory neurons follow more strongly and likely the magnitude of beta oscillations. Pairwise comparisons were performed using MWUT. *** p<0.001, **p<0.01, *p<0.05, whiskers of boxplots show the 5-95^th^ percentile.

**Supplementary Figure 2.**
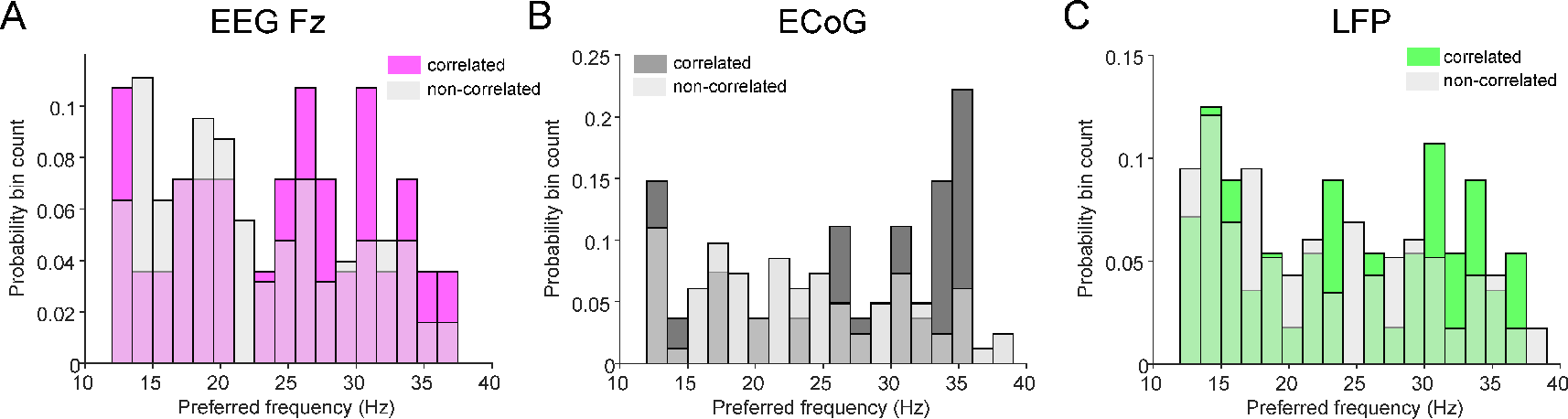
Preferred frequencies of phase-locking. **(A-C)**, Comparison of preferred frequencies of phase-locking between correlated and non-correlated (correlation between z-score vector length (phase-locking strength) and mean normalised magnitude in each percentile) phase-locking pairs for Fz (A), ECoG (B) and LFP (C). Note that only the frequency range from 12-40 Hz was considered and the preferred frequency was defined as the frequency with the lowest p-value of the Raileigh-test. Data show that the preferred frequency of phase-locking to beta oscillations is widely distributed and is not different between units that get entrained by the magnitude of network oscillations and those which do not (Fz: non-correlated units (n=126): 23.26 +/− 23.26 Hz, positively correlated (n=28): 24.31 +/− 7.43 Hz, MWUT *P*=0.43; ECoG: non-correlated units (n=82): 24.03 +/− 7.76 Hz, positively correlated (n=27): 26.69 +/− 8.53 Hz, MWUT *P*=0.11; LFP: non-correlated units (n=56): 22.76 +/− 7.8 Hz, positively correlated (n=53): 24.10 +/− 8.04 Hz, MWUT *P*=0.27).

**Supplementary Figure 3.**
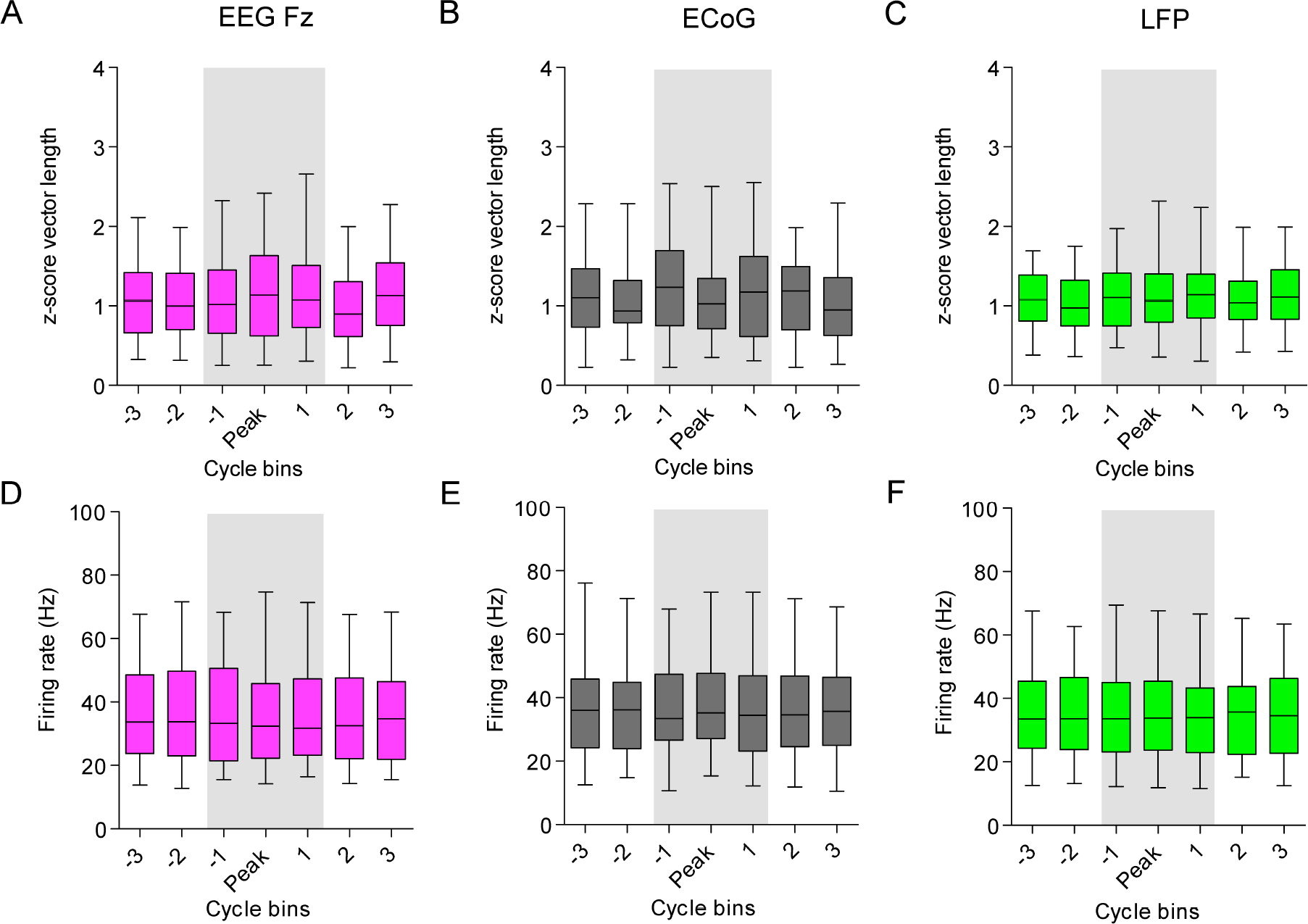
A subset of STN-neurons is not entrained during epochs elevated beta power (beta bursts) **(A-C)**, Beta burst analysis of STN-units, whose phase-locking strength is not significantly correlated with the magnitude of the ongoing network oscillation signal. Phase-locking analysis during episodes of elevated beta power detected with a threshold at the 75^th^ percentile of the magnitude for EEG Fz (A, pink, n=99), ECoG (B, grey, n=69) and LFP (C, green, n=94). Note that only STN unit-EEG/LFP pairs are shown which showed a negative or no correlation of phase-locking strength with the magnitude of the oscillation. X-axis showing the averaged phase-locking strength of spikes in each cycle bin. The grey shaded area shows the phase-locking during a beta burst aligned to the peak of the beta burst. Only beta bursts with a minimum duration of 3 cycles of the preferred beta frequency were included. In case of a longer burst duration, those cycle bins are not shown in the figure. The bins −2 and −3 show the phase-locking outside a beta burst with a distance of one cycle to the start of the beta burst, so that −2 is the second cycle bin to the edge of the burst and −3 the 3rd cycle bin before the start of the beta burst. Analogue the bins 2 and 3 show the 2nd and 3rd cycle bin after the end of the beta burst. **(D-E)**, Mean firing rate in corresponding cycle bins showing in A-C for STN units during episodes of elevated beta power in EEG Fz (D), ECoG (E) and LFP (F). A Kruskal-Wallis Anova reveals no significant difference for phase-locking strength and firing rate within and outside beta bursts. Therefor neither the phase-locking strength nor the firing rate for this subset of STN neurons was modulated during beta bursts.

**Supplementary Figure 4.**
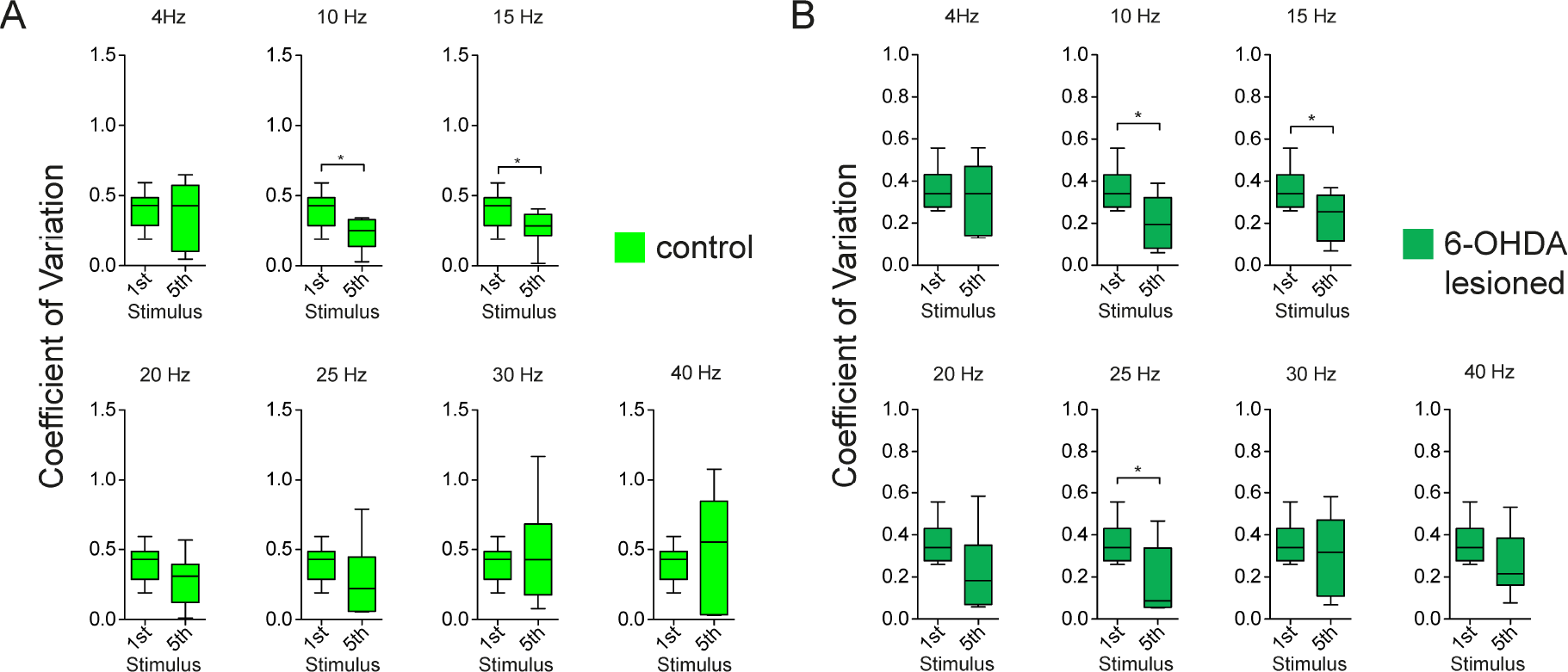
Similar pattern of entrainment of STN neurons to cortical stimulation in control and 6-OHDA lesioned rats. Comparison of the Coefficient of Variation (CV) of 1st and 5th stimulus (because the likelihood to detect a difference will increase with the course of stimuli) for control (n=8 STN neurons) **(A)** and 6-OHDA lesioned (n=8 STN neurons) **(B)** rats across frequencies. Box plots showing 5-95 percentile. * indicates p<0.05 of the signed Wilcoxon rank test were. Note the significant difference for 10 and 15 Hz in both groups, and the slightly more pronounced and significant difference at 25 Hz in 6-OHDA lesioned rats. Overall both groups show a trend of a lower CV at the 5th stimulus in the frequencies 10-25 Hz, indicating that the significant differences in the merged groups is caused by both groups.

**Supplementary Figure 5.**
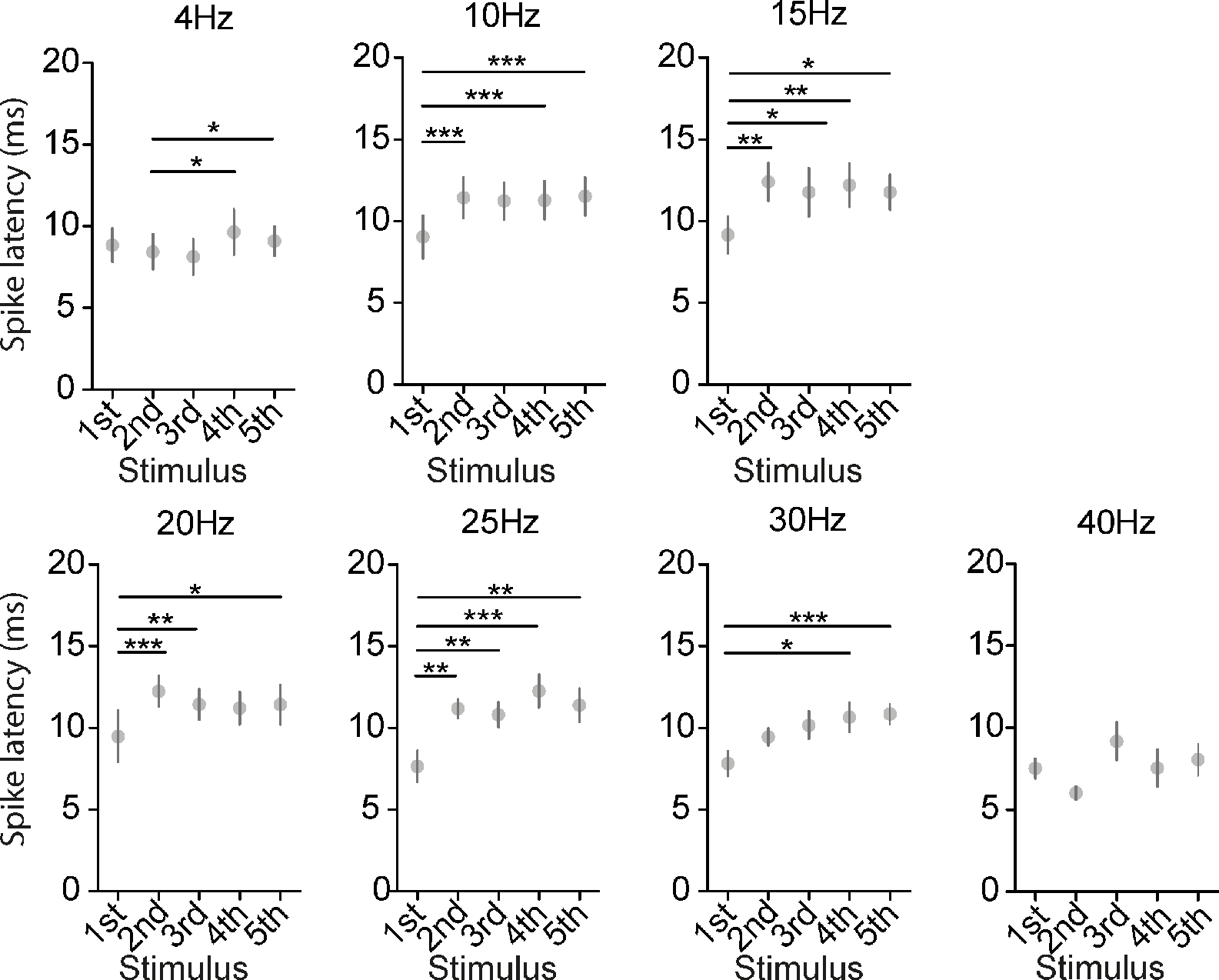
Increase of spike latency of the first spike over the course of cortical stimulation most pronounced at beta frequencies. Spike latency of the first spike after the stimulus in ms across stimuli and frequencies. Each subplot shows the firing probability over the course of stimuli at one frequency (at 4, 10, 15, 20, 25 and 30 Hz). Dots show the mean +/− SEM of the group (n=16 STN neurons). Note an increase spike latency mostly after the second spike for the frequencies 10,15,20 and 25 Hz. Overall, these results suggest an inhibition of spiking at the frequencies 10-30 Hz with a possible trend also for 4 Hz, which might even increase with additive stimuli. For comparison a Friedman test with post hoc Tukey comparisons was performed. *p<0.05, **p<0.01 and ***p<0.001

**Supplementary Figure 6.**
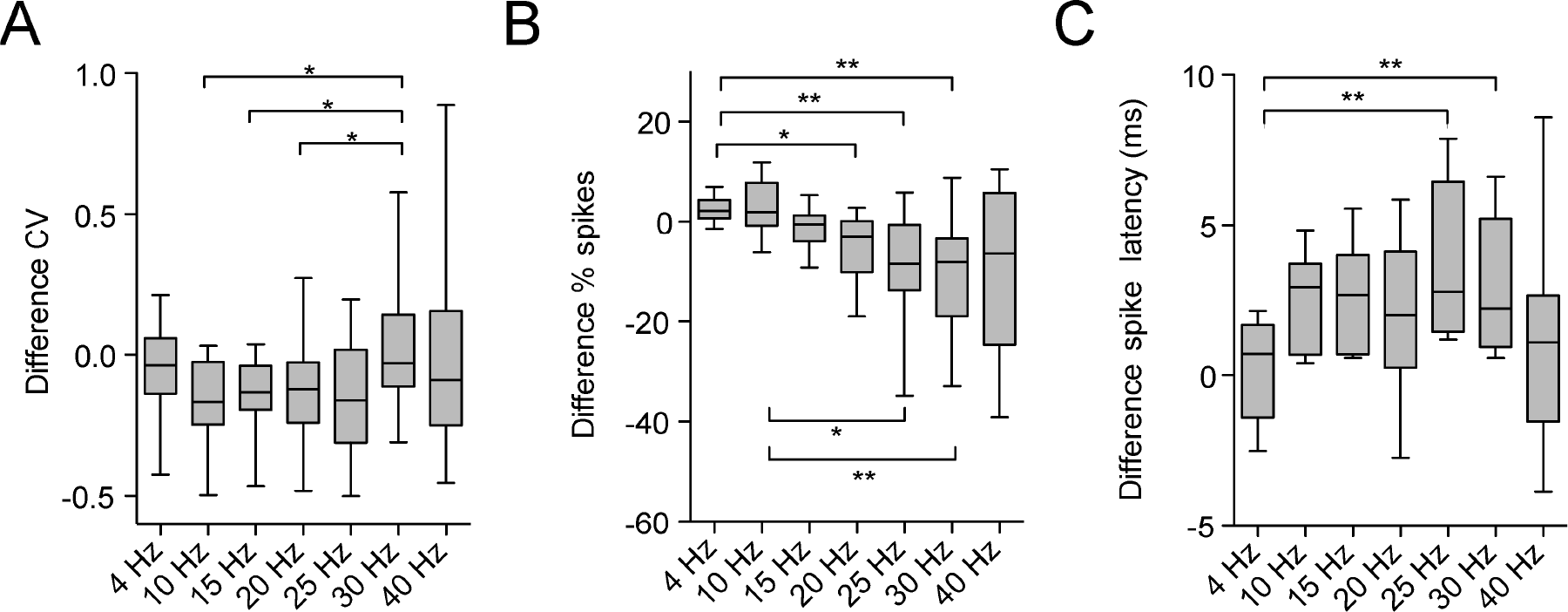
Entrainment characteristics of STN-neurons to cortical stimulation - comparison across frequencies. **(A-C)**, Difference between the 5th and 1st stimulus for the CV (A) (shown in Figure 8 across stimuli), firing probability in % of spikes (B) and spike latency to the stimulus (C) (n=16 STN neurons) (shown in Figure 9 across stimuli). Because the difference between 5th and 1st stimulus appears to be most accentuated, the difference was compared using a Friedman test with post-hoc Tukey’s comparison across frequencies. **(A)** Difference CV 5th stimulus-CV 1st stimulus. Negative values indicate that the CV of the first stimulus ist higher than the CV of the 5th stimulus. Note the tendency of a more frequent negative value in the frequencies 10-25 Hz in line with the statistic shown in Figure 8. **(B)** Difference % spikes after 5th stimulus - % spikes after 1st stimulus. Negative values indicate a decrease of firing probability after the 5th stimulus in comparison to the 1st stimulus. Note the decrease of firing probability over the course of stimuli with increasing frequency with an eminent difference between 4 Hz and 20, 25 and 30 Hz. Interestingly, in contrast to the CV comparison, 10 Hz is not behaving different from 4 Hz and has a significant higher difference as the frequencies 25 and 30 Hz. **(C)** Difference between the spike latency of the first spike to the 5th stimulus and 1st stimulus. A positive value indicates that the 5th stimulus has more frequently a longer spike latency in comparison to the 1st stimulus. Note the tendency of a longer spike latency after the 5th stimulus in the frequencies 10-30 Hz in comparison to 4 and 40 Hz with a significant difference between 4 Hz and 30 Hz. Overall these three metrics as a measurement of synchronization and rate coding underline a crucial difference in the behaviour of STN neurons to frequency selective response/entrainment to cortical stimulation in the frequencies 10-25 Hz. Whiskers of Boxplots show the 5-95th percentile. *p<0.05, **p<0.01 and ***p<0.001.

**Supplementary Table 1.** Patient details.

| Case | Age (Yrs) and Sex | Disease Duration (Yrs) | Motor UPDRS OFF | Motor UPDRS ON | Medication pre-operation | Hoehn/ Yahr Score | Dominant side | Major Symptoms | Hemispheres analysed | ECoG |
| --- | --- | --- | --- | --- | --- | --- | --- | --- | --- | --- |
| 1 | 61F | 25 | 50 | 30 | Levodopa 700 mg<br>Carbidopa 150 mg<br>Entacapone 1000 mg<br>Benserazide 25 mg<br>Pramipexol 0.26 mg | 5 | Left | Tremor,<br>Akinesia/rigidity<br>Fluctuations | Left | Yes |
| 2 | 70F | 8 | 38 | 21 | Ropinirole 8mg<br>Alpha-dihydroergocryptine 40 mg<br>Amantadine 600 mg<br>Levodopa 250mg<br>Carbidopa 150mg | 3 | Left | Bradykinesia<br>Fluctuations<br>Dyskinesia | Both | No |
| 3 | 67M | 25 | 55 | 19 | Levodopa 1250 mg<br>Entacapone 1400 mg<br>Carbidopa 312.5mg<br>Rotigotine 6 mg<br>Amatadine 150mg | 3 | Right | Equivalence<br>Fluctuations<br>Dyskinesia | Both | Yes |
| 4 | 64M | 15 | 53 | 39 | Amantadine 300 mg<br>Levodopa 550 mg<br>Ropinirole 20 mg<br>Entacapone 900 mg<br>Carbidopa 112.5 mg<br>Benserazide 25 mg | 4 | Left | Akinesia/rigidity<br>Camptocormia | Left | Yes |
| 5 | 69F | 9 | 21.5 | 7 | Levodopa 450 mg<br>Lisuride 0.9 mg<br>Rotigotine 4 mg<br>Amantadine 300 mg | 3 | Right | Equivalence type<br>Fluctuations<br>Dyskinesia | Both | Yes |
| 6 | 66M | 11 | 28 | 14 | Amantadine 150 mg<br>Levodopa 1450 mg<br>Tolcapone 300mg<br>Carbidopa 312.5<br>Benserazide 50mg | 4 | Right | Akinesia/rigidity<br>Fluctuations | Both | No |
| 7 | 72F | 18 | 41 | 19 | Levodopa 700mg<br>Carbidopa 175mg | 4 | Left | Equivalence type<br>Fluctuations<br>Dyskinesia | Left | Yes |
| 8 | 63F | 17 | 35 | 16 | Rotigotine 4mg<br>Ropinirole 25mg<br>Levodopa 350mg | 3 | Left | Akinesia/rigidity<br>Bradykinesia<br>Fluctuations<br>Dyskinesia | Both | Yes |
| 9 | 56M | 10 | 18 | 6 | Amantadine 300 mg<br>Safinamide 100 mg<br>Levodopa 450 mg<br>Benserazide 112.5<br>Opicapone 50 mg<br>Pramipexole 2.1 mg | 2 | Left | Equivalence type<br>Fluctuations | Both | No |
| 10 | 49M | 10 | 21 | 8 | Levodopa 725 mg<br>Entacapone 725 mg<br>Carbidopa 162.5 mg<br>Ropinirole 32 mg | 2 | Right | Bradykinesia<br>Akinesia/rigidity<br>Fluctuations | Both | No |
| 11 | 66M | 8 | 26 | 15 | Levodopa 300 mg<br>Benserazide 75 mg<br>Rasagiline 1 mg | 3 | Right | Bradykinesia<br>Akinesia/rigidity<br>Fluctuations<br>Camptocormia | Both | No |
| 12 | 74F | 13 | 32 | 22 | Amantadine 300 mg<br>Carbidopa 118.75 mg<br>Levodopa 575 mg<br>Rotigotine 14 mg | 2-3 | Left | Bradykinesia<br>Dyskinesia<br>Camptocormia | Both | Yes |
| 13 | 58M | 7 | 38 | 25 | Amantadine 200 mg<br>Benserazide 30.5 mg<br>Levodopa 550 mg<br>Pramipexole 3.15 mg | 2 | Right | Equivalence type<br>Fluctuations | Both | No |
| 14 | 53M | 10 | 46 | 11 | Benserazide 50 mg<br>Carbidopa 156.25 mg<br>Entacapone 725 mg<br>Levodopa 825 mg<br>Ropinirole 16 mg | 2-3 | Right | Equivalence type<br>Fluctuations | Both | Yes |
| 15 | 57M | 15 | 25 | 17 | Amantadine 300 mg<br>Benserazide 125 mg<br>Levodopa 650 mg<br>Pramipexole 2.1 mg<br>Tolcapone 300 mg | 2 | Left | Bradykinesia<br>Akinesia/rigidity<br>Fluctuations<br>Dysarthrophonia | Right | Yes |
| 16 | 57F | 13 | 36 | 15 | Amantadine 150 mg<br>Carbidopa 50 mg<br>Entacapone 200 mg<br>Levodopa 200 mg<br>Pramipexole 3.62 mg | 2 | Left | Bradykinesia<br>Akinesia/rigidity | Both | Yes |

**Supplementary Table 2.** Exact p-values of Tukey Kramer’s post-hoc comparison of the Coefficient of Variation of the first spike after each stimulus across consecutive stimuli within each frequency referring to Figure 9. The p-value for the Friedman test for each frequency is given and post hoc comparisons are shown only for significant Friedman tests with p<0.05.

|  |  |  |  |  |  |
| --- | --- | --- | --- | --- | --- |
| **4 Hz** Friedman test $P=0.12$ | | | | | |
| **10 Hz** Friedman test $P=0.0006$ | | | | | |
| **Stimulus** | **1** | **2** | **3** | **4** | **5** |
| **1** | N=16 | **P=0.001** | **P=0.02** | P=0.17 | **P=0.002** |
| **2** |  | N=16 | P=0.96 | P=0.52 | P=1 |
| **3** |  |  | N=16 | P=0.90 | P=1 |
| **4** |  |  |  | N=16 | P=0.59 |
| **5** |  |  |  |  | N=16 |
| **15 Hz** Friedman test $P=0.0005$ | | | | | |
| **Stimulus** | **1** | **2** | **3** | **4** | **5** |
| **1** | N=16 | **P=0.02** | **P=0.0002** | **P=0.02** | P=0.1 |
| **2** |  | N=16 | P=0.73 | P=1 | P=0.98 |
| **3** |  |  | N=16 | P=0.80 | P=0.38 |
| **4** |  |  |  | N=16 | P=0.96 |
| **5** |  |  |  |  | N=16 |
| **20 Hz** Friedman test $P=0.006$ | | | | | |
| **Stimulus** | **1** | **2** | **3** | **4** | **5** |
| **1** | N=16 | P=0.73 | **P=0.02** | **P=0.04** | **P=0.03** |
| **2** |  | N=16 | P=0.32 | P=0.52 | P=0.45 |
| **3** |  |  | N=16 | P=1 | P=1 |
| **4** |  |  |  | N=16 | P=1 |
| **5** |  |  |  |  | N=16 |
| **25 Hz** Friedman test $P=0.022$ | | | | | |
| **Stimulus** | **1** | **2** | **3** | **4** | **5** |
| **1** | N=16 | P=0.26 | P=0.13 | **P=0.02** | P=0.08 |
| **2** |  | N=16 | P=1 | P=0.80 | P=0.98 |
| **3** |  |  | N=16 | P=0.94 | P=1 |
| **4** |  |  |  | N=16 | P=0.98 |
| **5** |  |  |  |  | N=16 |
| **30 Hz** Friedman test $P=0.67$ | | | | | |
| **40 Hz** Friedman test $P=0.13$ | | | | | |

**Supplementary Table 3.** Exact p-values of Tukey Kramer’s post-hoc comparison of the % of spikes after each stimulus across consecutive stimuli within each frequency referring to Figure 10. The p-value for the Friedman test for each frequency is given and post hoc comparisons are shown only for significant Friedman tests with p<0.05.

|  |  |  |  |  |  |
| --- | --- | --- | --- | --- | --- |
| **4 Hz** Friedman test $P=0.003$ | | | | | |
| **Stimulus** | **1** | **2** | **3** | **4** | **5** |
| **1** | N=16 | **P=0.0006** | P=0.3 | P=0.36 | P=0.09 |
| **2** |  | N=16 | P=0.24 | P=0.19 | P=0.57 |
| **3** |  |  | N=16 | P=1 | P=0.98 |
| **4** |  |  |  | N=16 | P=0.96 |
| **5** |  |  |  |  | N=16 |
| **10 Hz** Friedman test $P=0.034$ | | | | | |
| **Stimulus** | **1** | **2** | **3** | **4** | **5** |
| **1** | N=16 | P=0.63 | P=0.88 | P=0.98 | P=0.48 |
| **2** |  | N=16 | P=0.13 | P=0.27 | **P=0.02** |
| **3** |  |  | N=16 | P=1 | P=0.96 |
| **4** |  |  |  | N=16 | P=0.83 |
| **5** |  |  |  |  | N=16 |
| **15 Hz** Friedman test $P=0.18$ | | | | | |
| **20 Hz** Friedman test $P=0.003$ | | | | | |
| **Stimulus** | **1** | **2** | **3** | **4** | **5** |
| **1** | N=16 | P=0.09 | P=0.07 | **P=0.002** | **P=0.02** |
| **2** |  | N=16 | P=1 | P=0.78 | P=0.98 |
| **3** |  |  | N=16 | P=0.78 | P=1 |
| **4** |  |  |  | N=16 | P=0.96 |
| **5** |  |  |  |  | N=16 |
| **25 Hz** Friedman test $P=0.0001$ | | | | | |
| **Stimulus** | **1** | **2** | **3** | **4** | **5** |
| **1** | N=16 | P=0.49 | **P=0.0004** | **P=0.005** | **P=0.005** |
| **2** |  | N=16 | P=0.09 | P=0.35 | P=0.35 |
| **3** |  |  | N=16 | P=0.96 | P=0.96 |
| **4** |  |  |  | N=16 | P=1 |
| **5** |  |  |  |  | N=16 |
| **30 Hz** Friedman test $P=0.002$ | | | | | |
| **Stimulus** | **1** | **2** | **3** | **4** | **5** |
| **1** | N=16 | P=0.93 | **P=0.045** | P=0.23 | **P=0.003** |
| **2** |  | N=16 | P=0.29 | P=0.71 | **P=0.045** |
| **3** |  |  | N=16 | P=0.96 | P=0.93 |
| **4** |  |  |  | N=16 | P=0.56 |
| **5** |  |  |  |  | N=16 |
| **40 Hz** Friedman test $P=0.15$ | | | | | |

**Supplementary Table 4.** Exact p-values of Tukey Kramer’s post-hoc comparison of the spike latency after each stimulus across consecutive stimuli within each frequency referring to supplementary Figure 4. The p-value for the Friedman test for each frequency is given and post hoc comparisons are shown only for significant Friedman tests with p<0.05.

| **4 Hz** Friedman test $P=0.004$ | | | | | |
| --- | --- | --- | --- | --- | --- |
| Stimulus | 1 | 2 | 3 | 4 | 5 |
| 1 | N=16 | P=0.67 | P=0.67 | P=0.38 | P=0.90 |
| 2 |  | N=16 | 1 | **P=0.02** | P=0.17 |
| 3 |  |  | N=16 | **P=0.02** | P=0.17 |
| 4 |  |  |  | N=16 | P=0.9 |
| 5 |  |  |  |  | N=16 |
| **10 Hz** Friedman test $P=3.69\text{e-}06$ | | | | | |
| Stimulus | 1 | 2 | 3 | 4 | 5 |
| 1 | N=16 | **P=2.62e-05** | **P=0.0005** | P=0.1 | **P=7.58e-05** |
| 2 |  | N=16 | P=0.96 | P=0.17 | P=0.1 |
| 3 |  |  | N=16 | P=0.52 | P=0.99 |
| 4 |  |  |  | N=16 | P=0.26 |
| 5 |  |  |  |  | N=16 |
| **15 Hz** Friedman test $P=0.0006$ | | | | | |
| Stimulus | 1 | 2 | 3 | 4 | 5 |
| 1 | N=16 | **P=0.001** | **P=0.04** | **P=0.001** | **P=0.02** |
| 2 |  | N=16 | P=0.85 | 1 | P=0.94 |
| 3 |  |  | N=16 | P=0.85 | P=1 |
| 4 |  |  |  | N=16 | P=0.94 |
| 5 |  |  |  |  | N=16 |
| **20 Hz** Friedman test $P=0.0003$ | | | | | |
| Stimulus | 1 | 2 | 3 | 4 | 5 |
| 1 | N=16 | **P=0.0005** | **P=0.001** | **P=0.1** | **P=0.02** |
| 2 |  | N=16 | P=0.1 | P=0.52 | P=0.90 |
| 3 |  |  | N=16 | P=0.67 | P=0.96 |
| 4 |  |  |  | N=16 | P=0.96 |
| 5 |  |  |  |  | N=16 |
| **25 Hz** Friedman test $P=8.58\text{e-}06$ | | | | | |
| Stimulus | 1 | 2 | 3 | 4 | 5 |
| 1 | N=16 | **P=0.007** | **P=0.007** | **P=2.69e-06** | **P=0.001** |
| 2 |  | N=16 | 1 | P=0.38 | P=0.99 |
| 3 |  |  | N=16 | P=0.38 | P=0.99 |
| 4 |  |  |  | N=16 | P=0.67 |
| 5 |  |  |  |  | N=16 |
| **30 Hz** Friedman test $P=0.0009$ | | | | | |
| Stimulus | 1 | 2 | 3 | 4 | 5 |
| 1 | N=16 | P=0.52 | **P=0.056** | **P=0.03** | **P=0.0005** |
| 2 |  | N=16 | P=0.8 | P=0.67 | P=0.1 |
| 3 |  |  | N=16 | P=0.1 | P=0.67 |
| 4 |  |  |  | N=16 | P=0.8 |
| 5 |  |  |  |  | N=16 |
| **40 Hz** Friedman test $P=0.08$ | | | | | |

